# Necrosis-induced apoptosis promotes regeneration in *Drosophila* wing imaginal discs

**DOI:** 10.1101/2021.04.01.438052

**Authors:** Jacob Klemm, Michael J. Stinchfield, Robin E. Harris

**Affiliations:** School of Life Sciences, Arizona State University, Tempe, AZ 85728

**Keywords:** regeneration, cell death, Drosophila, apoptosis, gene expression, necrosis, imaginal disc

## Abstract

Regeneration is a complex process that requires a coordinated genetic response to tissue loss. Signals from dying cells are crucial to this process and are best understood in the context of regeneration following programmed cell death, like apoptosis. Conversely, regeneration following unregulated forms of death such as necrosis have yet to be fully explored. Here we have developed a method to investigate regeneration following necrosis using the *Drosophila* wing imaginal disc. We show that necrosis stimulates regeneration at an equivalent level to that of apoptosis-mediated cell death and activates a similar response at the wound edge involving localized JNK signaling. Unexpectedly however, necrosis also results in significant apoptosis far from the site of ablation, which we have termed necrosis-induced apoptosis (NiA). This apoptosis occurs independent of changes at the wound edge and importantly does not rely on JNK signaling. Furthermore, we find that blocking NiA limits proliferation and subsequently inhibits regeneration, suggesting that tissues damaged by necrosis can activate programmed cell death at a distance from the injury to promote regeneration.

## Introduction

Apoptosis has a fundamental role in shaping organisms during development and in maintaining tissue homeostasis (Fuchs and Steller 2011; Suzanne and Steller 2013). This vital cellular process acts to counter infection, suppress tumor formation and promote recovery from damage (Fogarty and Bergmann 2017; Hacker 2018; Wong 2011). These events occur because dying cells release signals that act on surrounding cells, resulting in different outcomes depending on the nature of the signals and the tissue context (Fogarty and Bergmann 2015; Fuchs and Steller 2015; Perez-Garijo and Steller 2015; Su 2015). Numerous aspects of regeneration can be dependent on this apoptotic signaling, including morphogenesis, inflammation and tissue regrowth (Bergmann and Steller 2010; Perez-Garijo and Steller 2015; Vriz et al. 2014). In many organisms, factors released from dying cells stimulate the proliferation of adjacent cells as a replacement mechanism (Chera et al. 2009; Li et al. 2010; Pellettieri et al. 2010; Su 2015; Tseng et al. 2007; Vriz et al. 2014). This specialized form of compensatory proliferation is known as apoptosis-induced proliferation (AiP) (Fogarty and Bergmann 2017), in which a number of mitogens, including WNTs, TGFβ, EGF and Hedgehog, are released from dying cells (Fogarty and Bergmann 2017; Fuchs and Steller 2015). Upstream factors that lead to their production, such as JNK signaling and the activation of caspases, have also been described (Fogarty and Bergmann 2017; Pinal et al. 2019). Conversely, apoptosis can also promote the death of surrounding cells, a process known as apoptosis-induced apoptosis (AiA), (Fogarty and Bergmann 2015; Perez-Garijo et al. 2013; Su 2015). This process also relies on JNK signaling and possibly represents a mechanism to ensure the proper coordination of cell death across a tissue (Perez-Garijo et al. 2013). Signals from dying cells can also result in non-autonomous protection from cell death, ensuring the preservation of neighboring cells after injury (Bilak et al. 2014). Thus, cell death can strongly influence the behavior of surrounding cells and consequently regulate the ability of a tissue to recover from damage.

Evidence that signals from apoptotic cells impact surrounding tissues first originated from studies of the *Drosophila* wing imaginal disc, a larval epithelial tissue that forms the adult wing (Huh et al. 2004; Perez-Garijo et al. 2004; Ryoo et al. 2004). The wing disc has been extensively characterized as a model for growth and development, and for its significant capacity to regenerate (Beira and Paro 2016; Hariharan and Serras 2017; Worley et al. 2012). Much of our understanding of imaginal disc regeneration has derived from GAL4/UAS-based genetic ablation tools, in which tissue-specific apoptosis is induced by a temperature change, permitting the study of regeneration *in situ* (Fox et al. 2020). However, these ablation tools rely on activating the apoptotic machinery to induce damage, limiting our ability to explore other types of cell death. One example is necrosis, the rapid and disordered death of a cell characterized by the loss of membrane integrity and release of cytoplasmic contents into the surrounding tissue (Fuchs and Steller 2015). Several factors released by cells undergoing necrosis have been identified and characterized as signaling molecules that might influence the behavior of surviving tissues (Bianchi 2007; Patel 2018; Venereau et al. 2015), although their role in wound recovery and regeneration is currently not well understood. In humans, necrosis can occur in all types of tissues and results from environmental insults like burns or frostbite, microbial infection (Bonne and Kadri 2017; Hakkarainen et al. 2014), and from common ischemic events such as heart attacks and strokes (Konstantinidis et al. 2012; Radak et al. 2017; Yuan 2009). Various inherited and congenital conditions can also lead to necrosis, including joint disorders and sickle cell anemia (Masi et al. 2007; Milner et al. 1991). Thus, a better understanding of how a tissue responds to damage could advance therapeutic methods to improve healing and regeneration in many human disease and injury contexts.

Here we have developed a method to induce necrotic cell death in the wing imaginal disc and used it to characterize the genetic response to necrosis compared to apoptosis. Our results show that wing discs can regenerate following necrosis but do so by inducing extensive JNK-independent apoptosis at a distance from the site of injury. Importantly, we demonstrate that this necrosis-induced apoptosis is required for localized proliferation and efficient regeneration.

## Materials and Methods

### *Drosophila* stocks and crosses

Flies were cultured in conventional dextrose fly media at 25°C at 12 hr light-dark cycles. The recipe for dextrose media contains 9.3 g agar, 32 g yeast, 61 g cornmeal, 129 g dextrose, and 14 g tego in 1 L distilled water. Complete fly line details and RRIDs are available in the Reagent Table. Genotypes for each figure panel are listed in the Supplemental Methods File. The lines used as ablation stocks are as follows: *hs-flp; hs-p65; R85E08-DBD, DVE>>GAL4* (*DC^NA^), hs-flp ; lexAop-GluR1^Lc8^, hs-p65 ; salm-LexADBD, DVE>>GAL4 / S-T* (*DC^GluR1^), hs-flp ; lexAop-GluR1^Lc^, hs-p65 / CyO ; salm-LexADBD / TM6C, sb* (*DC^GluR1^ no GAL4*), *hs-flp ; lexAop-hep^CA^, hs-p65 ; R85E08-DBD, DVE>>GAL4/S-T* (*DC^hepCA^*), and *hs-flp ; lexaOp-rpr, hs-p65 / CyO, salm-LexADBD, DVE>>GAL4* (*DC^rpr^*,(Harris et al. 2020). The stock *DR^WNT^-GFP* was used as a marker for early regeneration (Harris et al. 2016). The following stocks were obtained from Bloomington Drosophila Stock Center: *UAS-y^RNAi^* (BL#64527), *UAS-p35* (BL#5073), *PCNA-GFP* (BL#25749), *UAS-bsk^RNAi^* (BL#53310), *UAS-bsk^DN^* (BL#9311), *UAS-egr^RNAi^* (BL#58993), *puc^A251^* (BL#11173), *hep^r75^* (BL#6761), *UAS-hepCA* (BL#58981), *UAS-wg^RNAi^* (BL#32994)*, UAS-Mkk4^RNAi^* (BL#35143), *UAS-hep^RNAi^* (BL#28710), *UAS-rpr^RNAi^* (51846), and *hid^1^* (BL#631). *CyclinE-GFP* was a gift from the Wei Du lab. *UAS-grnd^RNAi^* was a gift from the Bilder lab at UC Berkeley. *UAS-miRHG* and *UAS-nlsGFP; egr-GAL4* were gifts from the Hariharan lab at UC Berkeley. *UAS-GluR1^Lc8^* (Liu et al. 2014) was a gift from the Xie lab at Stowers Institute.

### Generation of lexAOp-hep^CA^, lexAOp-GluR1^Lc8^, and DR^WNT^-GAL80 lines

To generate *lexAop-hep^CA^*, the coding region for *hep^CA^* was cloned from *UAS-hep^CA^* stock and subcloned into the *lexAop-GFP* vector following excision of GFP, using Xhol and XbaI sites. *LexAop-GluR1^Lc8^* was generated by cloning out the *GluR1^Lc8^* coding region from *UAS-GluR1^Lc8^* and subcloning it into the *lexAop-GFP* vector following excision of the *GFP*, using *XhoI* and *Xbal* sites. The *lexAop-hep^CA^* and *lexAop-GluR1^Lc8^* constructs were sequence verified and inserted into the Su(Hw)attP5 landing site via PhiC31 insertion. To generate *DR^WNT^-GAL80*, the GAL80 coding sequence was cloned from tub-GAL80 genomic DNA (BL#5132) and subcloned into the *DR^WNT^-GFP* plasmid following excision of the *GFP* (Harris et al. 2016), using *AgeI* and *Xbal* sites. *DR^WNT^-GAL80* was sequence verified and inserted into the *attp40* and *VK00027* landing sites via PhiC31 insertion. Transgenic flies were generated by embryo injections performed by BestGene, Inc.

### Ablation Experiments

DUAL Control experiments were performed essentially as described in (Harris et al. 2020). Experimental crosses were cultured at 25°C and density controlled at 50 larvae per vial. Larvae were heat shocked at day 3.5 or 4.5 of development (84 and 108 hours after egg lay (AEL), respectively) by placing vials in a 37°C water bath for 45 minutes, followed by a return to 25°C. Larvae were allowed to recover for 18 hr before being dissected, fixed and stained, unless otherwise indicated. The non-ablating version of DUAL Control (*DC^NA^*) was used as a control while *UAS-y^RNAi^* was used as a control for RNAi experiments. DUAL Control experiments utilized the *DVE>>GAL4* driver that drives expression in the majority of the wing pouch (Figure S1A-B). In the experiment to examine *egr* (Figure S4H-J), *egr-GAL4* was utilized instead of *DVE>>GAL4*, which drives in the adult muscle precursor (AMP) cells of the disc (Everetts et al. 2021).

### Regeneration Scoring and Wing Measurements

Wings of adult flies from heat shocked larvae were scored and measured after genotype blinding by another researcher. Scoring was performed on anesthetized adults by binning into a regeneration scoring category (Figure 1G and (Harris et al. 2020). Wing measurements were performed by removing wings, mounting in Permount solution and imaging using a Zeiss Discovery.V8 microscope. Wing area was measured using the Image J software. Male and female adults were measured separately to account for sex differences in wing size, using a reproducible measuring protocol that excludes the variable hinge region of the wing (details of measuring protocol available on request). Statistics were performed using GraphPad Prism 9.0.

**Figure 1.**
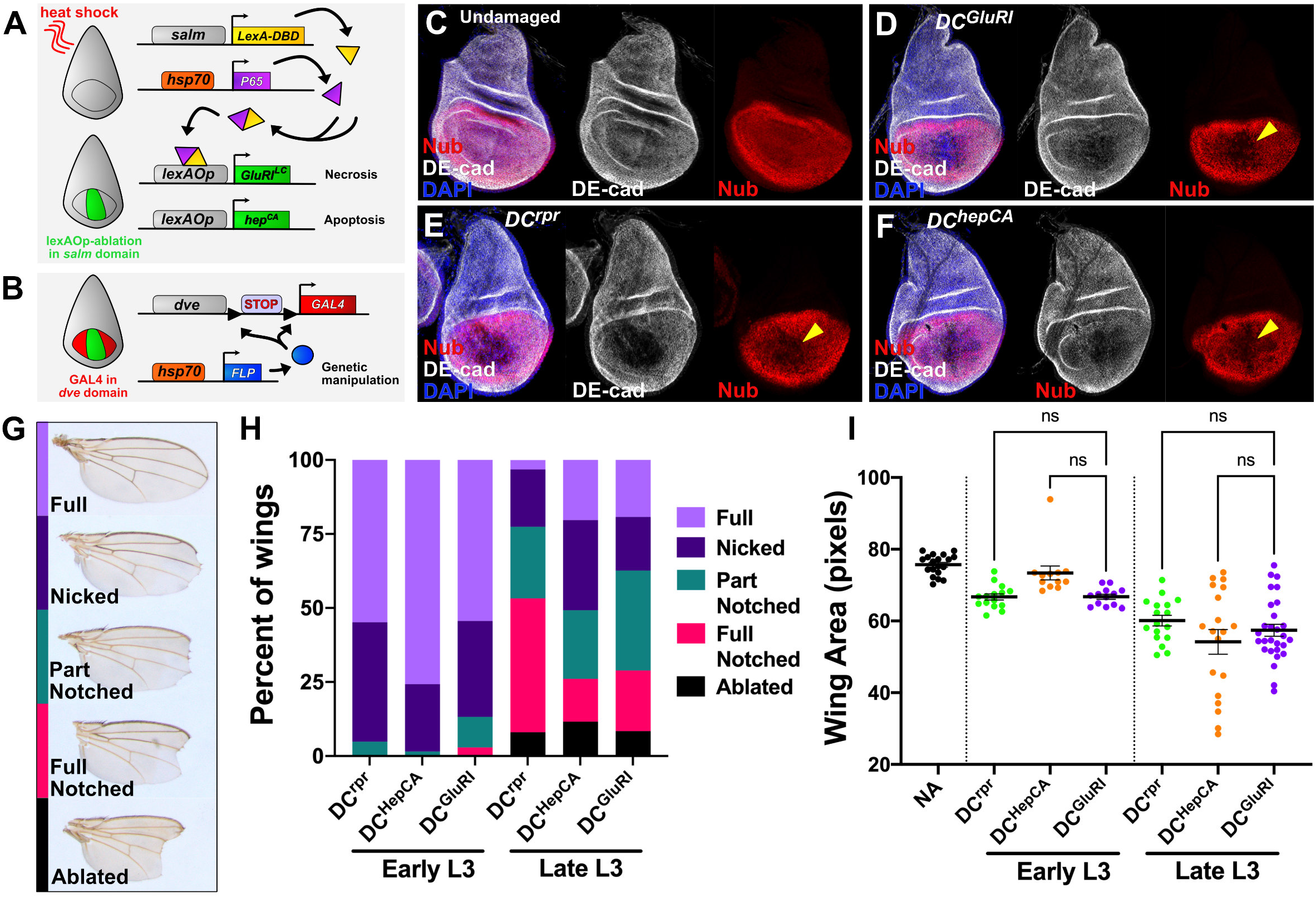
*Drosophila* wing imaginal discs regenerate comparably following ablation with *DC^rpr^*, *DC^hepCA^* and *DC^GluR1^.* (**A-B**) A schematic of the DUAL Control genetic ablation system. This system is based on a split LexA transcriptional activator which can form a complete transcriptional activator (yellow/purple protein) using complementary leucine zipper domains following a heat shock. This in turn induces *lexAOp*-driven ablation in the *salm* domain (green area of disc) (**A**). The *DVE>>GAL4* driver is used to drive *UAS-*based transgenes in the wing pouch independent of ablation (red area of disc), following the removal of a stop cassette by *hs-Flp* (**B**). Red lines in (**A**) indicate a heat shock. *salm*: *spalt* enhancer, *DBD: DNA Binding Domain*, *dve*: *defective proventriculus* enhancer, *Flp: Flp recombinase*. (**C-F**) Wing discs stained for the Nubbin pouch marker (Nub, red) and DE-cadherin (DE-cad, gray) in an unablated control (*DC^NA^)* (**C**) and following ablation with *DC^GluR1^* (**D**), *DC^rpr^* (**E**), and *DC^hepCA^* (**F**). DAPI, blue. Arrowheads in (**D**-**F**) indicate loss of Nub pouch marker. (**G**) Adult wing ablation phenotypes following DUAL Control ablation (see also Harris et al., 2020). (**H**) Regeneration scoring of adult wings from early L3 discs ablated by *DC^rpr^* (n = 62), *DC^hepCA^* (n = 66), and *DC^GluR1^* (n = 68), and late L3 discs ablated by *DC^rpr^* (n = 62), *DC^hepCA^* (n = 69), and *DC^GluR1^* (n = 83). (**I**) Mean pixel area of adult wings from early L3 ablated wings in an unablated control (NA, n = 19), *DC^rpr^* (n = 22), *DC^hepCA^* (n = 12), and *DC^GluR1^* (n = 17). ns, not significant. Data analyzed using one-factor ANOVA and Tukey’s multiple comparisons test.

### Immunohistochemistry

Larvae were dissected in PBS followed by a 20-minute fix in 4% paraformaldehyde in PBS. After 3 washes in 0.1% PBST (Triton-X), larvae were washed in 0.3% PBST and then blocked in 0.1% PBST with 5% NGS for 30 minutes. Primary staining was done overnight at 4°C, while secondary staining was done for 4 hours at room temperature. The following primary antibodies were obtained from the Developmental Studies Hybridoma Bank: mouse anti-Nubbin (1:25), mouse anti-Wg (1:100), mouse anti-Mmp1 C-terminus (1:100), mouse anti-Mmp1 catalytic domain (1:100), mouse anti-LacZ (1:100), and rat anti-DE-Cadherin (1:100). Rabbit anti-Dcp1 (1:1000) and mouse anti-PH3 (1:400) was obtained from Cell Signaling Technologies, and chick anti-GFP (1:2000) was obtained from Abcam. The secondary anti-Chick 488 antibody was also obtained from Abcam. Other secondary antibodies, anti-rabbit 647, anti-rat 647, and anti-mouse 555 were obtained from Invitrogen. All secondary fluorophore-conjugated antibodies were used at 1:500. Images were obtained on a Zeiss AxioImager.M2 with ApoTome. For each experiment at least 15 discs were analyzed prior to choosing a representative image. Images were processed using Affinity Photo and Affinity Designer.

### Propidium Iodide staining and TUNEL Assay

Propidium iodide (PI) staining was achieved by incubating freshly dissected wing imaginal discs in Schneider’s Media with 15 µM PI for 10 minutes, followed by several washes in PBS and immediately mounting and imaging. Samples were imaged on a Zeiss AxioImager.M2 with ApoTome. The TUNEL assay was performed with the ApopTag Red *In Situ* Apoptosis Detection Kit (Millipore, S7165). Dissected larvae were fixed in 4% paraformaldehyde, followed by a 10-minute wash in 75 µL equilibration buffer. Discs were then submerged in 55 µL working strength TdT enzyme for 3 hours at 37°C. The reaction was stopped by adding 1 mL stop/wash buffer and incubating for 10 minutes at room temperature, followed by 3 washes in PBS. Primary staining was done by incubating 65 µL of the anti-digoxygenin rhodamine overnight at room temperature.

### Quantification and Statistical Analysis

Adult wing measurements were gathered using the ImageJ software. GraphPad Prism 9.0 was used for statistical analysis and graphical representation. Graphs depict the mean of each treatment, while error bars indicate the 95% confidence interval. For quantification of DCP1 and PH3 positive cells, the cell counter plugin of ImageJ was used. For quantification of *PCNA-GFP* signal, mean fluorescence intensity in pixels was measured using ImageJ. To quantify Propidium Iodide (PI), due to the high variability of signal the area of PI detection was calculated as a percent of the area of the *lexAop-GFP* reporter. All experiments were repeated twice and the n for each experiment is indicated in the figure legends. Statistical significance was evaluated in Prism using one-sample T-test or One-Factor ANOVA, corrected for multiple comparisons by Tukey’s or FDR.

### Data and Reagent Availability

Strains and plasmids are available upon request, and details of stocks and reagents used in this study are available in the Reagent Table. The authors affirm that all data necessary for confirming the conclusions of the article are present within the article, figures, and tables.

## Results

### A genetic method to induce necrotic injury in the wing imaginal disc

To study the response to necrosis we developed a method to induce damage in the larval wing disc by adapting a system we previously established to study regeneration called DUAL Control (Harris et al. 2020). This system utilizes a split version of the bacterial *LexA*/*lexAOp* transcriptional regulator to transiently express a pro-apoptotic gene in the distal wing tissue under the control of the *spalt* (*salm*) enhancer following a short heat shock (Figure 1A). This apoptosis stimulates regeneration over the subsequent 24 to 48 hr, which can be quantitatively assessed by examining the degree to which the distal tip of the wing is present in the adult fly (Harris et al. 2020). We previously used *lexAOp*-driven *reaper* (*rpr*) to induce cell death (Harris et al. 2020), and have since established another apoptosis-promoting transgene, *lexAOp-hep^CA^*, a constitutive form of the JNK kinase *hemipterous* (Adachi-Yamada et al. 1999). To study necrosis, we used the GluR1 protein, a murine glutamate receptor that regulates Ca^2+^ ion entry into a cell (Kohda et al. 2000). A characterized allele of this ion channel (*GluR1^LC^*) confers constitutive activity, leading to Ca^2+^ influx, osmotic swelling and catastrophic death of the cell (Kohda et al. 2000; Yang et al. 2013). We generated *lexAOp-GluR1^LC^* and used it to build a version of DUAL Control that expresses *GluR1^LC^* in developing wing disc (Figure 1A and S1A-B).

To test our ability to induce damage using this system, we heat shocked larvae bearing DUAL Control expressing *GluRI^LC^* alongside discs ablated by *rpr* or *hep^CA^* (hereafter *DC^GluRI^*, *DC^rpr^* and *DC^hepCA^*), using the pouch marker *nubbin* (*nub*) and the structural marker *DE-cadherin* (*DE-cad*) to evaluate disc integrity (Figure 1C). Ablation with *DC^GluRI^* leads to a loss of tissue in the *salm* domain, shown by an absence of both markers (Figure 1D), which is similar to the loss of tissue resulting from ablation by *DC^rpr^* (Figure 1E) and *DC^hepCA^* (Figure 1F). Ablation only occurs when all three transgenes that comprise the DUAL Control system are present (Figure S1C-F). We next examined the degree to which wing discs can regenerate following DC^GluR1^ ablation compared to DC^rpr^ and DC^hepCA^. Regeneration was assayed by appearance of the adult wing, scored for the presence of the distal wing tip (Figure 1G and 1H) and measurement of the total wing area (Figure 1I). Ablation by *DC^rpr^* and *DC^hepCA^* in early L3 (84 hr after egg deposition, AED) leads to significant regeneration, illustrated by both wing scoring and area (Figure 1H-I). Ablation by *DC^GluRI^* results in similar levels of ablation and regeneration (Figure 1H-I). The ability of discs to regenerate is known to decline with developmental maturity from early to late L3 (Harris et al. 2016; Harris et al. 2020). Ablation by each method in late L3 (108 hr AED) results in a similar loss of distal wing tissue (Figure 1H-I). Together, these data show that *DC^GluRI^* can ablate the wing disc and induce regeneration at levels equivalent to that of apoptosis-based methods.

To characterize how *DC^GluRI^* induces damage we examined ablated discs for different cell death markers. The loss of tissue in *DC^GluRI^* coincides with strong propidium iodide (PI) staining (Figure 2A, arrowhead, S2A and S2C), which labels cells with compromised membranes such as those undergoing necrosis (Wlodkowic et al. 2011). Conversely, PI staining of *DC^hepCA^* ablated discs shows only weak PI (Figure 2B, S2B-C). As activated caspases are a hallmark of apoptosis but not necrosis, we also stained ablated discs for the cleaved caspase DCP1. As expected, ablation with *DC^rpr^* or *DC^hepCA^* yields a strong DCP1 signal concentrated within the *salm* domain (Figures 2E-F). Sporadic caspase-positive cells are seen outside of the ablation domain in all discs, likely due to the heat shock used to activate ablation (Figure 2C). Surprisingly, discs ablated with *DC^GluRI^* also have significant levels of DCP1 (Figure 2D). However, most of this staining is observed in the lateral wing pouch at a distance from the ablated tissue (Figure 2D, arrowheads). These DCP1 positive cells are clearly separable from the dying cells in the *salm* domain. They are also not a result of dying cells moving away from the site of injury, as a *lexAop-GFP* transgene that labels ablated cells fails to mark the caspase positive cells outside of the *salm* domain (Figures 2G and 2H, arrowheads, S2D, S2F). This contrasts with *DC^hepCA^* discs where GFP and DCP1 almost entirely overlap (Figure 2I and 2J, S2E), with very few DCP1-positive cells observed outside of the *salm* domain (Figure S2F). Thus, the observed apoptosis does not occur in cells killed by *GluRI* but is instead induced in neighboring cells.

**Figure 2:**
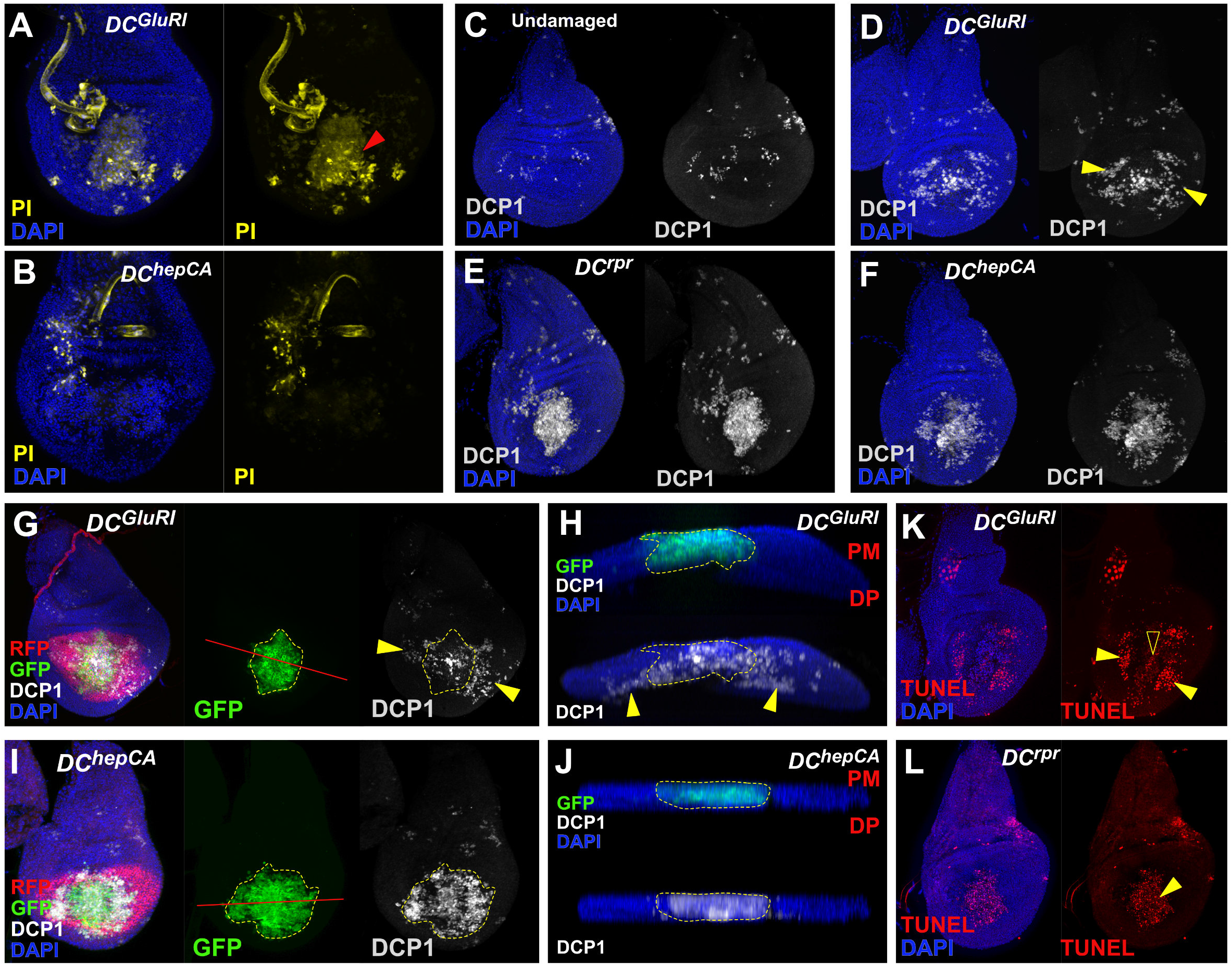
Ablation with *DC^GluR1^* generates apoptosis at a distance. (**A-B**) Propidium iodide (PI) staining (yellow) of *DC^GluR1^* (**A**) and *DC^hepCA^* (**B**) ablated wing discs. Arrowhead in (**A**) indicates PI-stained dying cells in the *salm* domain of the pouch. Non-specific PI staining of tracheal tubes is observed in both samples. DAPI, blue. (**C-F**) Caspase staining (DCP1, gray) of wing imaginal discs in an unablated control (**C**), and following ablation with *DC^GluR1^* (**D**), *DC^rpr^* (**E**), and *DC^hepCA^* (**F**). DAPI, blue. Arrowheads in (**D**) indicate non-autonomous caspase outside of the ablation domain. (**G-J**) Wing imaginal discs bearing the *lexAop-GFP; UAS-RFP* reporter and ablated with *DC^GluR1^* (**G-H**) or *DC^hepCA^* (**I-J**). GFP (green) labels the *salm* ablation domain, RFP (red) labels the surrounding unablated pouch, and DCP1 (gray) labels apoptotic cells. DAPI, blue. (**G**) Discs ablated with *DC^GluR1^* shows DCP1-positive cells outside of the ablation domain (arrowheads), red line indicates transverse section shown in (**H**). (**H**) A transverse section through the pouch of the disc in (**G**) confirms that these lateral DCP1-positive cells are not labeled with GFP. (**I**) Ablation by *DC^hepCA^* shows almost complete overlap between the GFP and DCP1 signals, red line indicates transverse section shown in (**J**). (**J**) A transverse section of the disc in (**I**) confirming significant overlap of GFP and DCP1. PM: peripodial membrane, DP: disc proper. (**K-L**) TUNEL-labeling (red) of wing discs ablated with *DC^GluR1^* (**K**) and *DC^rpr^* (**L**). Solid arrowheads indicate strong TUNEL staining in both panels, open arrowhead in (**K**) indicates weak TUNEL staining present in the *salm* domain.

In addition to promoting apoptosis, caspases can perform other cellular functions in imaginal discs unrelated to cell death (Su 2020). To examine whether the detected caspase activity is related to cell death we also performed a TUNEL assay, which reveals strong staining in the caspase positive cells found outside of the *salm* domain in *DC^GluRI^* discs (Figure 2K, arrowheads). This is consistent with *GluRI*-induced caspase activity leading to cell death, as in *DC^rpr^* ablated discs (Figure 2L). We also observed a separate weaker TUNEL signal in *DC^GluRI^* ablated discs within the *salm* domain, indicative of DNA fragmentation (Figure 2K, open arrowhead), which has previously been described in necrotic cells (Grasl-Kraupp et al. 1995). Together, these results demonstrate that ablation with *DC^GluR1^* leads to tissue loss and stimulates regeneration, but also unexpectedly activates widespread apoptosis in the surrounding cells at a distance from the injury.

### Tissue loss via *DC^GluRI^* ablation occurs independent of caspase activity

To understand if the non-autonomous apoptosis observed in *DC^GluRI^* discs contributes to tissue loss, or alternatively is a consequence of it, we used the baculoviral caspase inhibitor P35 to block apoptosis in the entire pouch (Hay et al. 1994). DUAL Control permits the expression of *UAS*-driven transgenes in the wing pouch independent of ablation under the control of a flip-out GAL4 (*DVE>>GAL4*), which is initiated by the same heat shock that triggers ablation and persists throughout the subsequent regeneration period (Figure 1B and S1A-B (Harris et al. 2020)). To test P35 in the DUAL Control system, we first blocked apoptosis in *rpr* ablated discs using *UAS-p35* (*DC^rpr^>>p35*). These discs have an intact pouch epithelium, shown by Nub and DE-cad staining (Figure 3A), and lack TUNEL staining (Figure S3A). DCP1 is still detected in *DC^rpr^>>p35* discs although the intense punctate staining is replaced by a diffuse cytoplasmic signal (Figures 3C and 3D), which has been reported previously when caspase activity is inhibited by P35 (Perez-Garijo et al. 2013). Conversely, P35 is unable to prevent tissue loss resulting from *DC^GluRI^* ablation, indicated by the absence of Nub and DE-cad in the pouch (Figure 3B) and presence PI staining in the *salm* domain (Figure 3G). The appearance of the DCP1 staining in *DC^GluRI^>>p35* discs is also diffuse (Figures 3E and 3F) like that of *DC^rpr^>>p35*, while TUNEL staining in the lateral pouch is now absent (Figure 3H), indicating that the apoptosis in these regions is being blocked. However, the weak TUNEL signal within the cells of the *salm* domain is still observed (Figure 3H, open arrowhead). These data suggest that inhibition of caspase activity cannot block tissue loss caused by *DC^GluR1^* ablation.

**Figure 3.**
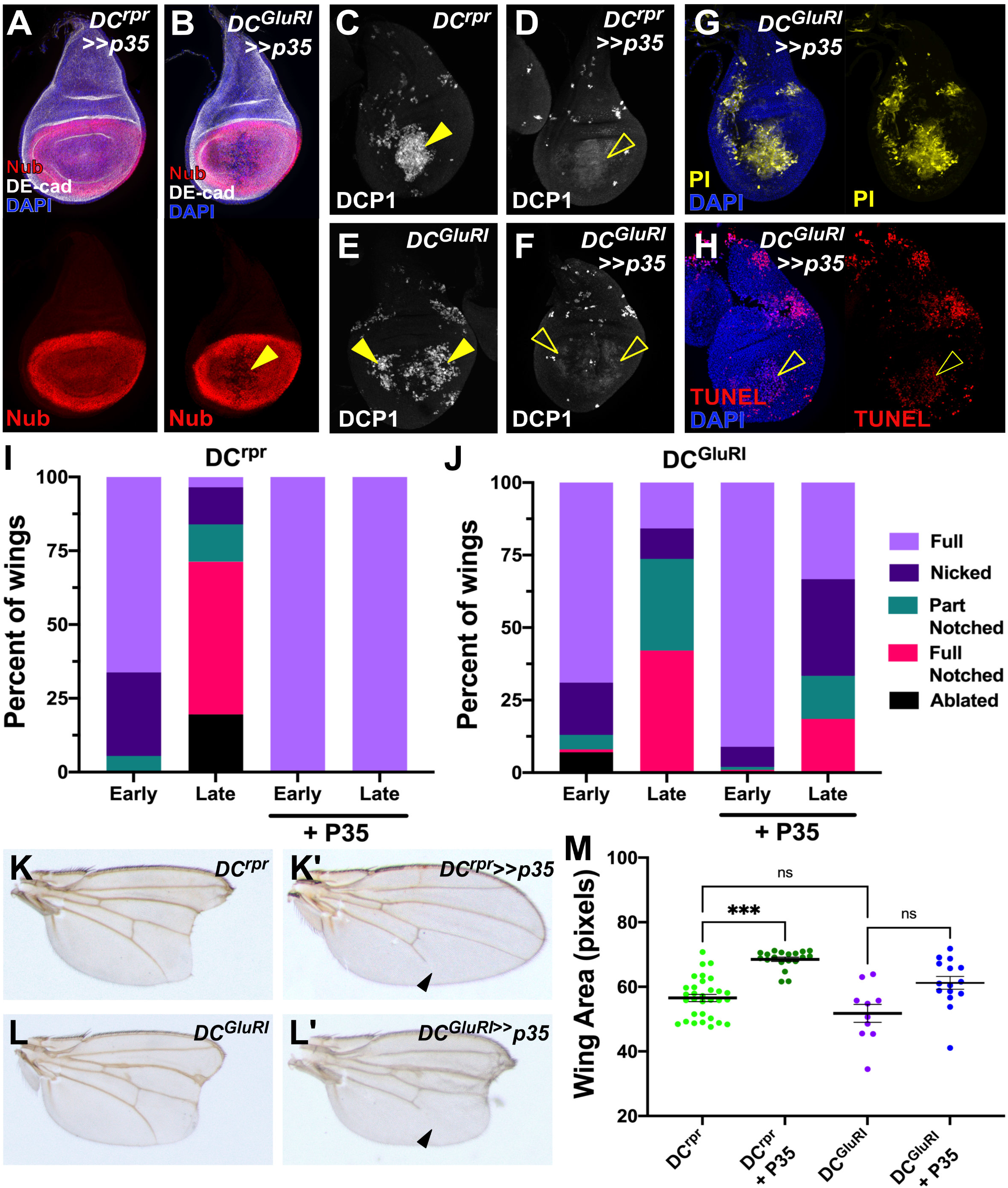
Inhibition of caspase activity fails to prevent ablation in DC^GluR1^ discs. (**A-B**) Wing discs stained for Nub (red) and DE-cad (gray) following ablation with *DC^rpr^>>p35* (**A**) and *DC^GluR1^>>p35* (**B**). DAPI, blue. Loss of tissue by *DC^GluRI^* ablation in the presence of P35 is indicated by the arrowhead in (**B**). (**C-F**) DCP1 staining (gray) in wing discs ablated with *DC^rpr^* (**C**), *DC^rpr^>>p35* (**D**), *DC^GluR1^* (**E**), *DC^GluR1^>>p35* (**F**). The arrowheads indicate strong DCP1 staining in the ablated *salm* domain in (**C**) and non-autonomous apoptosis in (**E**), while the open arrowheads in (**B**) and (**F**) indicate the diffuse, cytoplasmic DCP1 staining in these populations that occurs in the presence of P35. (**G**) PI staining (yellow) of a *DC^GluR1^>>p35* ablated wing disc. Non-specific PI staining of the tracheal tube is observed. DAPI, blue. (**H**) TUNEL-labeling (red) of a *DC^GluR1^>>p35* ablated wing disc. DAPI, blue. The open arrowhead indicates weak TUNEL staining in the *salm* domain, while the lateral TUNEL staining is no longer observed in the presence of P35. (**I-J**) Regeneration scoring of adult wings from early and late L3 discs ablated by *DC^rpr^* (**I**) or *DC^GluR1^* (**J**) in the presence and absence of P35. Number of wings scored: early and late L3 *DC^rpr^* (n = 74, n = 87), early and late L3 *DC^rpr^>>p35* (n = 110, n = 125), early and late L3 *DC^GluRI^* (n = 100, n = 38), early and late L3 *DC^GluRI^>>p35* (n = 101, n = 27). (**K-L’**) Adult wing phenotypes following ablation with *DC^rpr^* (**K**), *DC^rpr^>>p35* (**K’**), *DC^GluR1^* (**L**), and *DC^GluR1^>>p35* (**L’**). Arrowheads in (**K’**) and (**L’**) indicate an incomplete L5 vein, which is a consequence of *p35* expression on vein development, reported previously (Perez-Garijo et al. 2004). (**M**) The mean area of adult wings from discs ablated with *DC^rpr^* (n = 31), *DC^rpr^>>p35* (n = 19), *DC^GluR^*^1^ (n = 10), and *DC^GluR1^>>p35* (n = 15), ns, not significant, *** p = 0.0002, data analyzed by one-factor ANOVA and Tukey’s multiple comparisons test.

In agreement with the observed disc phenotypes, blocking apoptosis with P35 in *rpr* ablated discs results in full sized adult wings, shown by scoring and wing area measurements of both early and late L3 ablated organisms (Figures 3I, 3K-K’ and 3M). By contrast, blocking apoptosis in *DC^GluRI^* ablated discs does not prevent tissue loss, as adult wings that develop from *DC^GluRI^>>p35* discs have similar phenotypes to discs ablated in the absence of P35 (Figures 3J, 3L-L’ and 3M). Together these observations suggest that *DC^GluR1^* can kill cells independent of caspase activity, and that the apoptotic cells seen at a distance from the wound are a consequence of this tissue loss rather than being required for it. Thus, we refer to *DC^GluRI^*-induced tissue loss as necrotic ablation, and the non-autonomous caspase-positive cell death in the surrounding tissue as necrosis-induced apoptosis (NiA).

In our adult wing assays we noted a distinct category of wings from *DC^GluRI^>>p35* animals that are poorly patterned and have increased blistering and folding (Figure S3B and S3C). This phenotype also occurs in *DC^hepCA^*>>*p35* discs (Figure S3C). Blocking caspase activity with P35 in cells undergoing apoptosis can produce undead cells that continue to express mitogenic genes like *wg* and *dpp* downstream of JNK signaling, leading to tumorous overgrowth of the surrounding tissue (Martin et al. 2009; Perez-Garijo et al. 2004; Ryoo et al. 2004). Indeed, *DC^hepCA^*>>*p35* have significant ectopic *wg* expression in the ablation domain overlapping the diffuse DCP1 compared to ablation in the absence of P35 (Figure S3D and S3E). We hypothesized that blocking apoptosis in *DC^GluRI^>>p35* ablated discs might result in undead cells that produce the overgrown adult wing phenotype. However, in *DC^GluRI^>>p35* discs ectopic *wg* is only detected near to the wound edge, and not in the surrounding pouch where undead NiA would be expected to produce ectopic *wg* (Figure S3F), and *Dpp* expression shown by a *Dpp-lacZ* reporter is not altered (data not shown). To test whether the observed *wg* is important for overgrowth we knocked it down in the entire pouch (*DC^GluRI^>>p35, wg^RNAi^),* which eliminated the overgrowth phenotype (Fig S3G-I). Since ectopic *wg* is only associated with apoptotic cells at the wound edge it is possible that it is only these cells that contribute to overgrowth when forced to become undead. To test this, we used a *DR^WNT^-GAL80* transgene in *DC^GluRI^>>p35* discs to limit p35 expression to the regions of the pouch where NiA cells occur (Figure S3J-L). In this experiment the overgrowth phenotype is significantly reduced (Figure S3C), demonstrating that NiA do not strongly contribute to overgrowth after becoming undead. Alongside the observation that they do not express *wg*, these data suggest that undead NiA cells do not activate the same genetic pathways as undead cells generated by directly activating apoptosis in the presence of P35.

### NiA and wound edge apoptosis originate separately

To investigate the origin and timing of the NiA seen in *DC^GluRI^* discs we performed a time course of ablation and regeneration (Figure 4). To observe the formation of regenerating cells we took advantage of the *DR^WNT^* transgene, a damage-specific fluorescent reporter of WNT expression, which is activated upon ablation and represents one of the earliest signaling events in disc regeneration (Figure 4A-4F)(Harris et al. 2016). *DR^WNT^* expression becomes apparent at the wound edge 6 hr post-ablation with *DC^GluR1^* (Figure 4B), while most apoptotic cells in this region become obvious at around 12 hr (Figure 4C, arrowhead). NiA populations are still not observed at this timepoint and only occur at 18 hr post ablation after the appearance of wound edge apoptosis and *DR^WNT^* activity (Figure 4D, arrowheads). The NiA persists as regeneration proceeds (Figure 4E), and by 48 hr, when *DR^WNT^* expression is diminished, the DCP1-positive cellular debris appears to be pushed dorsally and ventrally possibly as a result of pouch growth (Figure 4F). At this time point the pouch is fully restored, indicated by *nub* expression (Figure 4G). Interestingly, *DR^WNT^* activity is never observed beyond the wound edge and does not occur in NiA cells. Thus, the time course shows that formation of NiA is not only spatially distinct from the wound edge apoptosis and damage associated signaling shown by *DR^WNT^*, but is also temporally separated from it. These observations suggest that these two populations might arise via different mechanisms.

**Figure 4.**
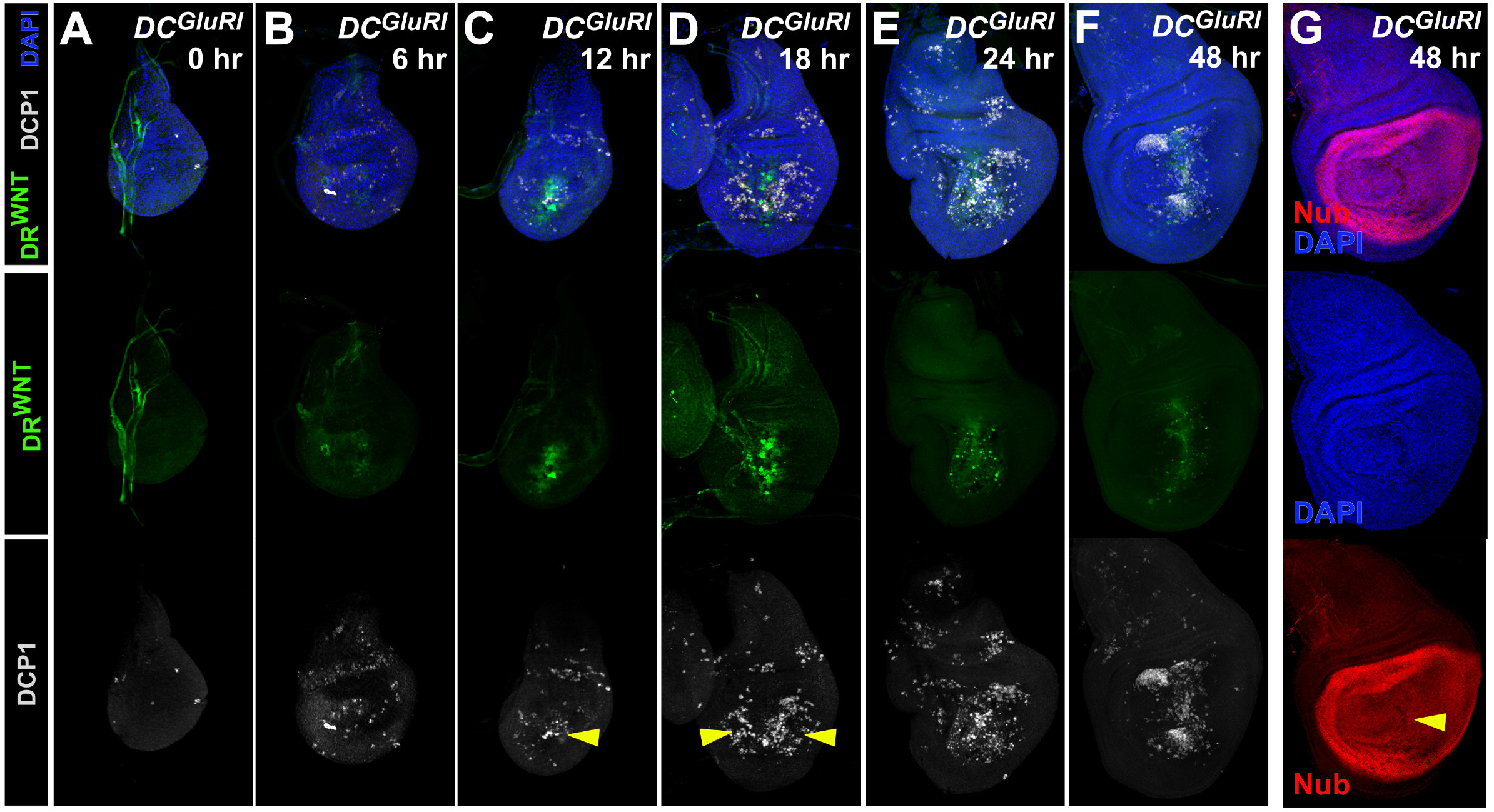
Necrosis-induced apoptosis (NiA) forms after wound edge signaling occurs. (**A-F**) A time course of regeneration for wing discs bearing a fluorescent reporter of a damage-responsive WNT enhancer (*DR^WNT^*, green, (Harris et al. 2016) ablated with *DC^GluR1^* and stained for DCP1 (gray) at (**A**) 0 hr, (**B**) 6 hr, (**C**) 12 hr, (**D**) 18 hr, (**E**) 24 hr, (**F**) 48 hr after heat shock. DAPI, blue. Arrowhead in (**C**) indicates wound edge apoptosis at 6 hr coinciding with the appearance of *DR^WNT^* expression, arrowheads in (**D**) indicates NiA at 18 hr. (**G**) A wing imaginal disc at 48 hr of recovery following ablation with *DC^GluR1^*. The pouch marker Nub (red) and DAPI staining (blue) show continuity of the pouch is restored (arrowhead).

### NiA is independent of JNK signaling

The *DR^WNT^* reporter used in the time course is directly activated by JNK signaling (Harris et al. 2016). Since *DR^WNT^* expression is restricted to the wound edge, and JNK signaling is a key pathway that regulates both apoptosis and proliferation in response to damage (Pinal et al. 2019), we evaluated JNK signaling in *DC^GluRI^* ablated discs. Using a reporter for AP-1 activity (*AP-1-GFP* (Chatterjee and Bohmann 2012)) we found that JNK signaling is present only in wound edge cells and not in cells undergoing NiA (Figure 5A), consistent with the *DR^WNT^* reporter expression. Examination of the JNK target genes *puc, wg* and *Mmp1* supports these results (Figures 5B, 5C and 5D), suggesting that necrotic ablation leads to JNK upregulation and low levels of apoptosis at the wound edge, and induces NiA at a distance without activating JNK.

**Figure 5.**
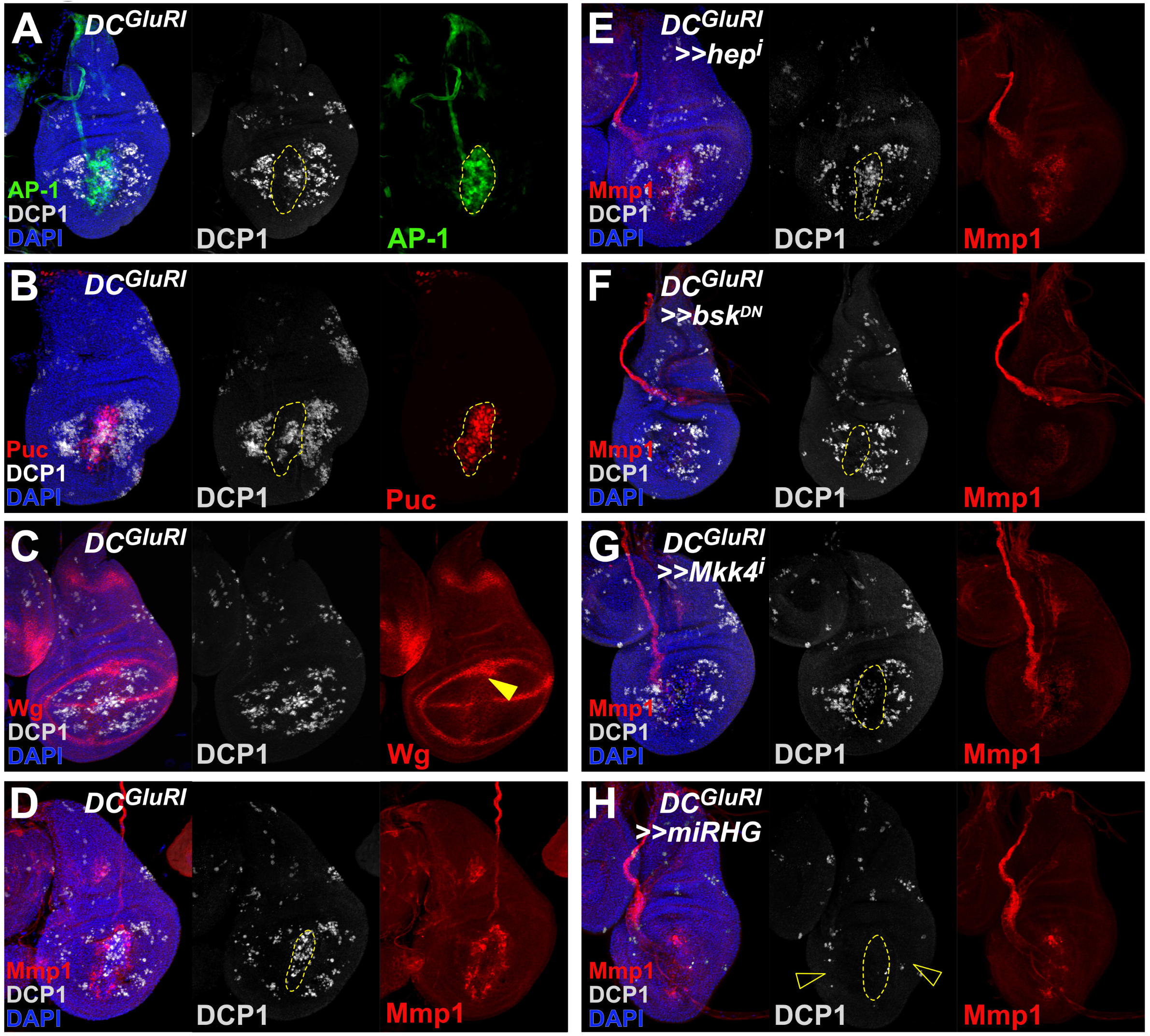
Necrosis-induced apoptosis (NiA) acts independent of the JNK pathway. (**A-B**) Wing discs ablated with *DC^GluR1^* and bearing the *AP-1-GFP* (green) (**A**) and *puc-lacZ* (red) (**B**) reporters of JNK pathway activity, stained for DCP1 (gray). DAPI, blue. These reporters show strong overlap with the wound edge apoptosis, but not with the NiA cells. As in other panels, the dashed line outlines the ablated *salm* domain, indicated by the appearance of cells labeled by DAPI. (**C-D**) Wing imaginal discs ablated with *DC^GluR1^* and stained for the JNK targets Wg (red) (**C**) and Mmp1 (red) (**D**). Arrowhead in (**C**) indicates ectopic Wg at the wound edge. DCP1, gray; DAPI, blue. (**E**) A wing imaginal disc ablated with *DC^GluR1^>>hep^RNAi^* stained for DCP1 (gray) and Mmp1 (red), DAPI, blue. Both the wound edge apoptosis and NiA cells remain unaffected following knockdown of *hep*. (**F**) A wing imaginal disc ablated with *DC^GluR1^>>bsk^DN^* stained for DCP1 (gray) and Mmp1 (red). DAPI, blue. Wound edge apoptosis is largely absent, while NiA cells remain unaffected. (**G**) A wing imaginal disc ablated with *DC^GluR1^>>Mkk4^RNAi^* stained for DCP1 (gray) and Mmp1 (red). DAPI, blue. Wound edge apoptosis is strongly reduced, while NiA cells remain unaffected. (**H**) A wing imaginal disc ablated with *DC^GluR1^>>miRHG* stained for DCP1 (gray) and Mmp1 (red). DAPI, blue. Both the wound edge apoptosis and NiA (open arrowheads) are eliminated following knockdown of *rpr, hid,* and *grim* with *miRHG*.

To test whether NiA does indeed occur independent of JNK, we investigated how altering different elements of the JNK pathway might influence either the wound edge or NiA populations in *DC^GluRI^* ablated discs. Knockdown of the JNK kinase *hep* using RNAi reduces the expression of the JNK target *Mmp1* but does not strongly alter apoptosis at the wound edge (Figure 5E). Importantly, NiA cells are still observed in the absence of *hep* (Figure 5E). This result is confirmed using a *hep* hemizygous mutant (*hep^r75^/Y*), although the level of apoptosis across the whole disc is somewhat reduced when using this allele (Figure S4A). The expression of a dominant negative form of the JNK *basket* (*UAS-bsk^DN^*) strongly suppresses *Mmp1* expression (Figure 5F), and unlike the loss of *hep*, *bsk^DN^* also reduces apoptosis at the wound edge (Figure 5F), suggesting that this cell death is dependent on *bsk*. The population of NiA cells is also unaffected in this background (Figure 5F), supporting the idea that NiA formation does not depend on JNK signaling. RNAi knockdown of *bsk* also confirms these results (Figure S4B). Since the loss of *bsk*, but not *hep*, seems to influence wound edge apoptosis, we also targeted an alternative JNK kinase, *Mkk4*, for knockdown in *DC^GluRI^* ablated discs. Unlike *hep*, reduction of *Mkk4* levels reduces both the apoptosis and *Mmp1* expression at the wound edge, but again has no influence on the appearance of NiA (Figure 5G). We also reduced expression of a TNF receptor *grindelwald* (*grnd*) and TNF ligand *egr* using RNAi. Neither manipulation affected the apoptosis or JNK activity at the wound edge or the appearance of the cells undergoing NiA outside the ablated domain (Figure S4C and S4D). Moreover, a reporter for *egr* expression (*egr-GAL4 ; UAS-GFP*) shows no upregulation of *egr* at the wound edge or in cells undergoing NiA (Figure S4H-S4J). Interestingly however, *egr* expression that is normally observed in adult muscle precursor cells under the notum is reduced upon ablation (Figure S4H-S4J). Overall, these data suggest that necrotic ablation induces low levels of apoptosis and JNK signaling at the wound edge, which occurs downstream of a JNK receptor *grnd* but is dependent on both *Mkk4* and *bsk*, while also generating NiA at a distance via a JNK-independent signal.

As NiA does not appear to require JNK, we sought to investigate factors upstream of the observed activated caspase DCP1 that might be required to produce NiA following necrotic ablation. We used an RNAi targeting the dIAP1 inhibitor *rpr*, which activates cell death immediately upstream of initiator caspases. Knockdown of *rpr* leads to a moderate reduction in apoptosis at the wound edge, but also reduces NiA formation (Figure S4E). Mutation of another dIAP1 inhibitor, *head involution defective* (*hid*), did not have an effect (Figure S4F). However, when a miRNA targeting *rpr*, *hid* and another dIAP1 inhibitor *grim* was used (*UAS-miRHG*), NiA formation was eliminated (Figure 5H), as was the cell death normally observed at the wound edge, indicating that more than one of these genes are required for both wound edge apoptosis and NiA. Importantly, *DC^GluRI^*-induced tissue loss still clearly occurs even in the absence of NiA (Figure S4G), while JNK signaling indicated by *Mmp1* expression is observed, albeit at a reduced level (Figure 5H). JNK signaling is thought to activate a feedforward mechanism through DRONC, reactive oxygen species (ROS) and hemocytes (Amcheslavsky et al. 2018), and therefore this diminished Mmp1 labeling could be due to interference with this mechanism. Overall, these results suggest that necrotic ablation leads to apoptosis at the wound edge and NiA in the surrounding pouch, and although each is regulated by different activating signals, they rely on the same downstream effectors, *rpr, hid,* and *grim*, to kill cells.

### NiA is required for damage-induced proliferation and subsequent regeneration

To understand the potential contribution of NiA to regeneration we examined how the appearance of NiA coincides with regenerative proliferation following necrotic ablation. We used reporters for *E2F* (*PCNA-GFP*) and *Cyclin E* (*cycE-GFP*), and PH3 staining to examine changes in proliferation that occur during regeneration. These markers show that proliferation induced by necrotic ablation does not significantly increase in the wing pouch until around 24 hr, after the formation of NiA at 18 hr (Figure 6A-B, S5A-E). To test whether NiA regulates this localized proliferation, we sought to eliminate this apoptosis following *DC^GluRI^* ablation. As NiA is unaffected by blocking JNK signaling, and P35 allows dying cells to persist in the disc which could complicate our analysis due to the observed overgrowth (Figure S3), we used *UAS-miRHG* to prevent the formation of NiA. The lack of NiA in the pouch following necrotic ablation results in a loss of *PCNA-GFP* upregulation that is normally seen at 24 hr (Figure 6C), as well as diminished PH3 levels (Figure S5E), suggesting that NiA is required for regenerative proliferation. Importantly, JNK signaling at the wound edge is still observed following *miRHG* expression, indicated by the presence of *Mmp1* (Figure 6C), demonstrating that proliferation following necrotic ablation is not dependent solely on JNK activity from the wound, but also requires NiA.

**Figure 6.**
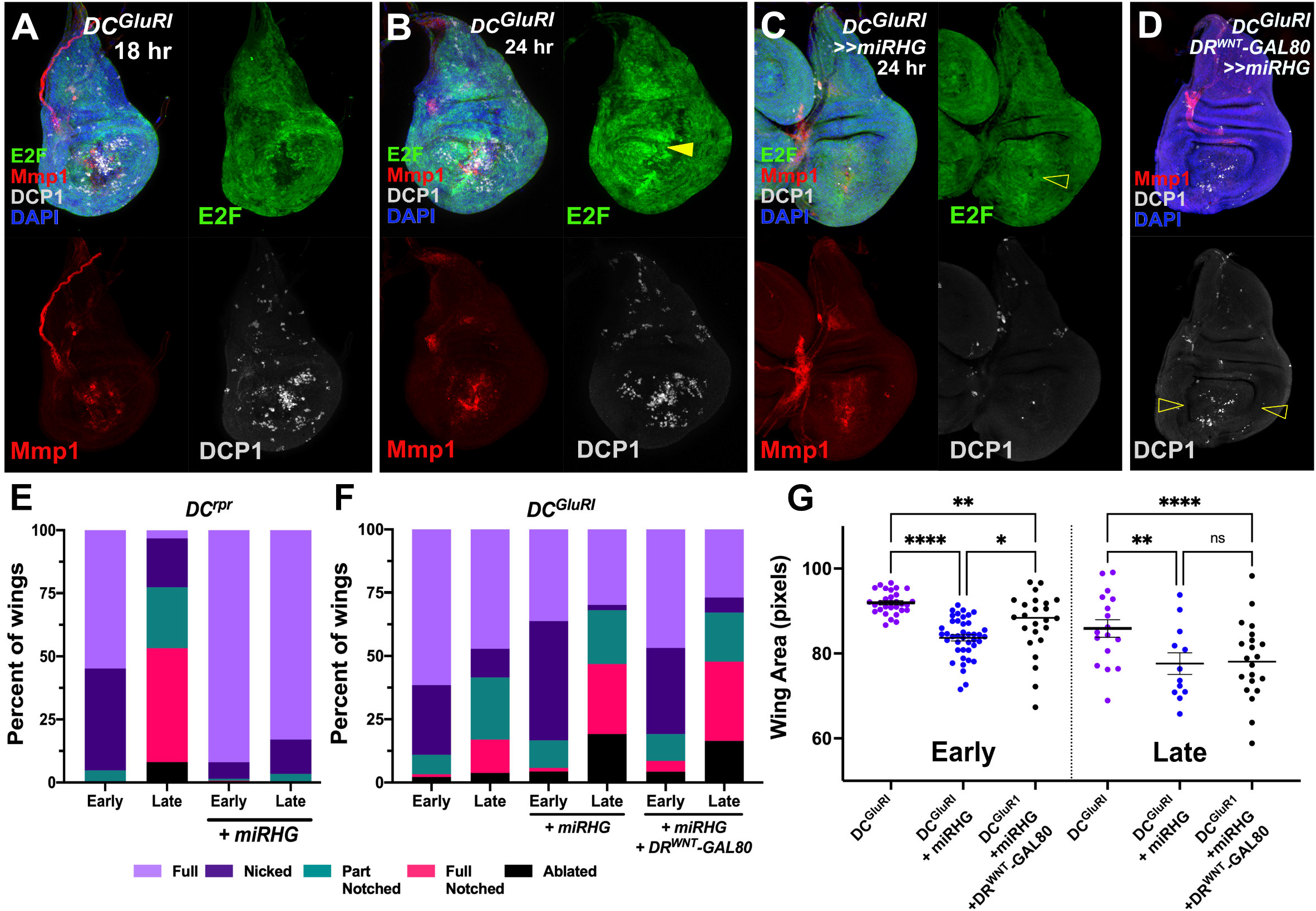
Blocking NiA formation impairs regeneration. (**A-B**) Wing discs bearing an *E2F* transcriptional reporter (*PCNA-GFP,* green) ablated with *DC^GluR1^* and stained for Mmp1 (red) and DCP1 (gray) at 18 hr (**A**) and 24 hr (**B**) of recovery. DAPI, blue. Arrowhead in (**B**) shows the localized proliferation that is only present after 24 hr of recovery. (**C**) Wing disc bearing *PCNA-GFP* (green) ablated with *DC^GluR1^>>miRHG* and stained for Mmp1 (red) and DCP1 (gray), DAPI, blue. The open arrowhead indicates reduced proliferation upon *miRHG* expression. JNK signaling, indicated by Mmp1 staining, is still present in these discs. (**D**) A wing imaginal disc ablated by *DC^GluR1^>>miRHG* in the presence of *DR^WNT^-GAL80* and stained for DCP1 (gray) and Mmp1 (red). DAPI, blue. Open arrowheads indicate a loss of NiA. (**E-F**) Regeneration scoring of adult wings from early and late L3 discs ablated with *DC^rpr^* (**E**) or *DC^GluR1^* (**F**) in the presence and absence of *miRHG*. Number of wings scored: early and late L3 *DC^rpr^* (n = 62, n = 63), early and late L3 *DC^rpr^>>miRHG* (n = 137, n = 88), early and late L3 *DC^GluRI^* (n = 91, n = 53), early and late L3 *DC^Glu1r^>>miRHG* (n = 138, n = 47), and early and late L3 *DC^GluR1^>> miRHG, DR^WNT^-GAL80* (n = 47, n = 67). While *miRHG* leads to more full and nicked wings following *DC^rpr^* ablation, the opposite is seen following *DC^GluR1^* ablation and *DC^GluR1^*+*DR^WNT^-GAL80*. (**G**) Mean pixel area of adult wings from early and late L3 discs ablated with *DC^GluR1^* (early n = 30, late n = 17), *DC^GluR1^>>miRHG* (early n = 41, late n = 12), and *DC^GluR1^>> miRHG, DR^WNT^-GAL80* (early n = 24, late n = 22). ns: not significant, ** p = 0.0056 (early *DC^GluRI^* vs. *DC^GluRI^ + miRHG + DR^WNT^-GAL80*), **** p <0.0001, * p =0.0254, ** p = 0.0015 (late *DC^GluRI^* vs. *DC^GluRI^ + miRHG*), data analyzed by one-factor ANOVA and FDR’s multiple comparisons tests.

We next examined how blocking NiA might influence the discs capacity to regenerate. Evaluating adult wings that develop from *DC^GluRI^>>miRHG* discs shows they have a reduced ability to regenerate compared to control discs (Figure 6F-G). This result is significant, as blocking apoptosis with *miRHG* would normally be expected to improve regeneration, as it does in *DC^rpr^* ablated discs (Figure 6E). However, in this experiment apoptosis is blocked throughout the pouch and is not specific to the NiA. Therefore, we used *DR^WNT^-GAL80* to limit *miRHG* expression solely to the surrounding pouch where NiA develops and allow wound edge apoptosis to still occur (Figure 6D). When wound edge apoptosis can still occur but NiA is blocked, regeneration is still strongly inhibited in both early and late L3 discs (Figure 6F-G), demonstrating that NiA is specifically required for regeneration in response to necrosis.

## Discussion

### Necrosis induces apoptosis at a distance in the wing imaginal disc

Necrotic ablation induces apoptosis in cells in two regions of the injured disc: at the wound edge and more extensively in the surrounding pouch. These two populations of dying cells are likely induced by separate activating signals, rely on the same downstream effectors, and contribute differently to regeneration (Figure 7). The wound edge apoptosis coincides with JNK activity and is likely dependent on it, as interference with *bsk* strongly limits the caspase observed at the wound. The factors that can activate JNK signaling, and those downstream that regulate whether cells subsequently proliferate or undergo apoptosis, are complex and context dependent (La Marca and Richardson 2020; Pinal et al. 2019). However, our experiments show that pathway activation following necrotic ablation does not rely on a receptor for JNK, *grnd*, and requires the JNK kinase *Mkk4* more than *hep*. There are a number of signals that can induce JNK activity in this way, including loss of polarity, mechanical inputs and the release of immediate stress signals such as ROS (La Marca and Richardson 2020). ROS has been shown to upregulate both JNK and P38, another factor that regulates proliferation and apoptosis, via Ask1/AKT (Santabarbara-Ruiz et al. 2019; Santabarbara-Ruiz et al. 2015), while ROS is known to be an important regulator of regeneration in the wing disc (Brock et al. 2017; Khan et al. 2017; Santabarbara-Ruiz et al. 2019; Santabarbara-Ruiz et al. 2015). Thus, it is possible that JNK signaling and the associated apoptosis at the wound edge induced by necrotic ablation is mediated via this pathway. It is likely that this JNK activity contributes to recovery following necrosis, as it does in apoptosis-induced regeneration. However, our observation that blocking NiA limits regeneration, despite wound edge apoptosis and JNK being unaffected, suggests that regeneration does not solely depend on JNK signaling.

**Figure 7.**
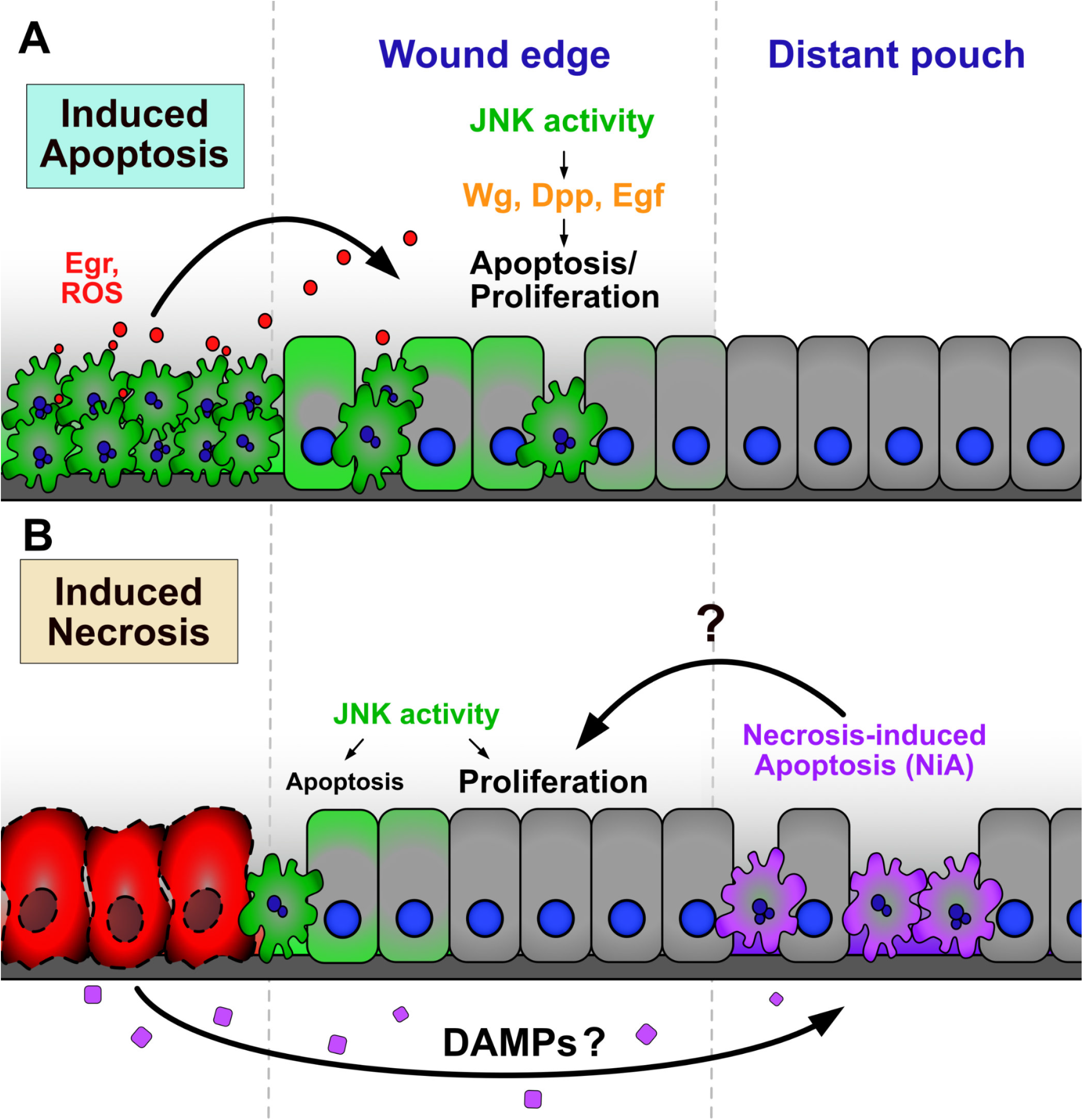
NiA cells promotes regeneration in *DC^GluR1^* ablated discs. (**A-B**) Schematics depicting the cell signaling events that occur resulting from apoptotic ablation (**A**) and necrotic ablation (**B**). Previous studies have shown dying cells release reactive oxygen species (ROS) and promote Egr release via hemocytes and potentially other mechanisms, inducing high levels of JNK signaling at the wound edge (Fogarty and Bergmann 2017). In turn, wound edge apoptotic cells and surviving cells with lower JNK signaling release mitogens including Wg, Dpp, and Egf, to promote regenerative signaling in the adjacent wound edge tissue. In cells undergoing necrosis, low levels of JNK signaling occurs downstream of the receptor at the wound edge, likely through pathways related to polarity disruption, mechanical stress, or immediate stress signals such as ROS. This signaling induces sporadic apoptosis at the wound edge that potentially contributes to proliferation and recovery. Necrotic cells also release an as yet unidentified long-range signal that could be a damage associated molecular pattern (DAMP), resulting in the formation of NiA cells in the distant pouch. These cells promote proliferation formation and regeneration.

The signals that lead to the NiA at a distance from the necrotic injury are less clear, however. A related phenomenon to the one we have observed, in which cells undergoing programmed cell death induce additional non-autonomous apoptosis, has been described in the wing imaginal disc and the mouse hair follicle (Perez-Garijo et al. 2013). However, in both contexts these events are induced by apoptotic cell death rather than necrosis, and are mediated by JNK signaling, unlike NiA. In a parallel observation to that of apoptotic cells communicating with surrounding tissues, studies have shown that cells undergoing necrosis might also release factors that regulate the behavior of neighboring cells. These factors are collectively known as <u>d</u>amage <u>a</u>ssociated <u>m</u>olecular <u>p</u>atterns or DAMPs (Gong et al. 2020; Roh and Sohn 2018). Such signals often consist of endogenous intracellular molecules like cytoskeletal components, DNA, RNA, histones and metabolites, which are passively released and inappropriately exposed to the extracellular environment. DAMPs are detected by immune cells that in turn activate immune pathways to mount a stress response known as sterile inflammation (Gong et al. 2020). Beyond these initial inflammatory responses, DAMPs are also important for orchestrating the subsequent steps in tissue repair and regeneration (Venereau et al. 2015). In *Drosophila*, the actin-binding protein α-actinin has been shown to act as a DAMP when purified and injected into adult flies, inducing a humoral immune response in the form of cytokine signal activation specifically in the fat body (Gordon et al. 2018; Srinivasan et al. 2016). Similarly, apoptotic-deficient *Drosophila* constitutively activate immune signaling in the larval fat body due to the presence of circulating DAMPs in the hemolymph (Ming et al. 2014). Recently it was shown that imaginal disc damage leads to the upregulation of circulating metabolites produced by the fat body, which are necessary for proper disc repair (Kashio and Miura 2020), highlighting the importance of interorgan communication in the context of regeneration. Thus, one possibility is that necrotic cells release DAMP signals to induce pro-regenerative NiA in the disc via an immune response involving the fat body. The role of hemocytes, an essential population of immune cells, in the regeneration of damaged imaginal discs has also been explored following physical and apoptotic damage (Fogarty et al. 2016; Katsuyama and Paro 2013; Pastor-Pareja et al. 2008), and in the context of sterile inflammation that occurs following necrosis (Shaukat et al. 2015). Considering the vital roles hemocytes play in these processes it is possible they also mediate immune or other signals to induce NiA formation. Alternatively, DAMPs are also known to have stimulatory effects on non-immune cells (Chen and Nunez 2010; Gong et al. 2020), and thus could be sensed directly by the immediately surrounding pouch tissues to induce NiA. Going forward it will be important to ascertain both the identity of the signal and the physiological mechanism by which necrosis leads to non-autonomous cell death in the wing disc.

### NiA is necessary for proliferation and subsequent regeneration

Our data show that blocking NiA following necrosis inhibits the localized proliferation that promotes tissue repair, strongly suggesting that NiA is a vital part of the recovery process. How this type of cell death might promote proliferation is still unclear, however. Apoptosis-induced proliferation (AiP) is a well-established phenomenon, in which dying cells release mitogens, such as *wg* and *dpp*, downstream of JNK signaling to stimulate localized proliferation (Fogarty and Bergmann 2017; Perez-Garijo and Steller 2015). These mitogens act via a feed forward mechanism that involves the release of ROS, recruitment of hemocytes and induction of further JNK signaling to promote repair (Amcheslavsky et al. 2018; Fogarty and Bergmann 2017; Fogarty et al. 2016). By contrast, we have shown that cells undergoing NiA do not detectably express *wg* or *dpp*, even in an undead state, and do not activate JNK signaling. Although we have not ruled out other characterized mitogens being involved, these findings suggest that NiA may not promote proliferation via the established AiP mechanism. An important consideration is whether cells undergoing NiA release specific signals to direct tissue regrowth, as seen with AiP, or if the tissue loss resulting from necrosis and NiA is instead sensed more generally at the level of the whole disc, stimulating coordinated regrowth in addition to localized proliferation (Hariharan 2015). The identification of signals produced by NiA cells and a better understanding of proliferation rates across a recovering disc will help to reveal the contribution of these different mechanisms to necrosis-induced regeneration.

Tissue necrosis is associated with many types of traumatic injury and diseases in humans, including ischemic injuries, infections and cancer. Using model systems, like the one we have developed here, to understand how necrotic cells impact neighboring healthy tissues, and how these tissues enact mechanisms to effectively recover, may help to identify therapeutic interventions that promote healing in tissues which otherwise struggle to recover from necrosis, such as those of humans.

## Supporting information

Regents table

## Acknowledgements

The authors would like to thank Dr. David Bilder and Dr. Iswar Hariharan of UC Berkeley and Dr. Tian Xie of the Stowers Institute for their gift of stocks. We thank Ayla Zustra and current members of the Harris lab for useful feedback. We thank the Bloomington Stock Center, Developmental Studies Hybridoma Bank, and BestGene for stocks, reagents, and services.

## Funding Information

The funders had no role in study design, data collection and interpretation, or the decision to submit the work for publication. This work was supported by a grant from the NICHD 1 R21 HD102765-01 to Robin Harris.

## Competing interests

No competing interests declared.

## Supplemental Figure Legends

**Figure S1. Related to Figure 1.**
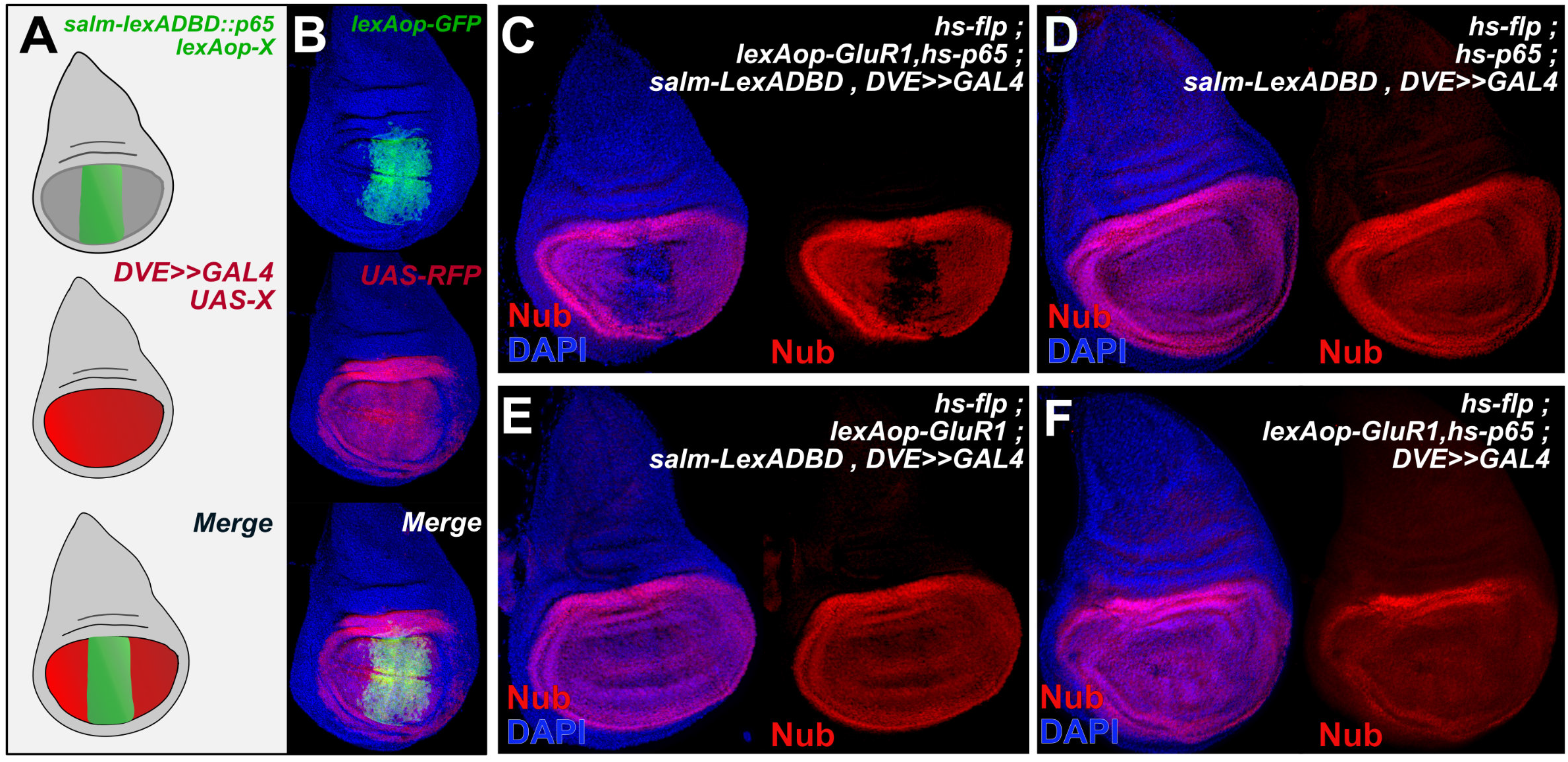
Ablation and gene expression using DUAL Control. (**A**) A schematic of the domains used for ablation and genetic manipulation in most DUAL Control experiments. The ablation domain coincides with an enhancer of the *salm* gene, which patterns the distal wing pouch. The domain used for genetic manipulation utilizes an enhancer of the *dve* gene, which drives *UAS*-based constructs in the entire wing pouch. (**B**) A wing imaginal disc bearing *DC^NA^* with *lexAop-GFP* to mark the ablation domain (green) and *UAS-RFP* to mark the pouch domain (red), as diagrammed in the schematic in (**A**). DAPI, blue. (**C-F**) Wing discs bearing combinations of the transgenes that comprise DUAL Control, showing that heat shock induced ablation only occurs when all three genes (*hsp65, lexAop-GluRI, salm-LexADBD*) are present, as indicated by the presence of Nub (red) in the *salm* domain. DAPI, blue. Genes omitted: (**D**) *lexAop-GluR1* (**E**) *hs-p65* (**F**) *salm-LexADBD*.

**Figure S2. Related to Figure 2.**
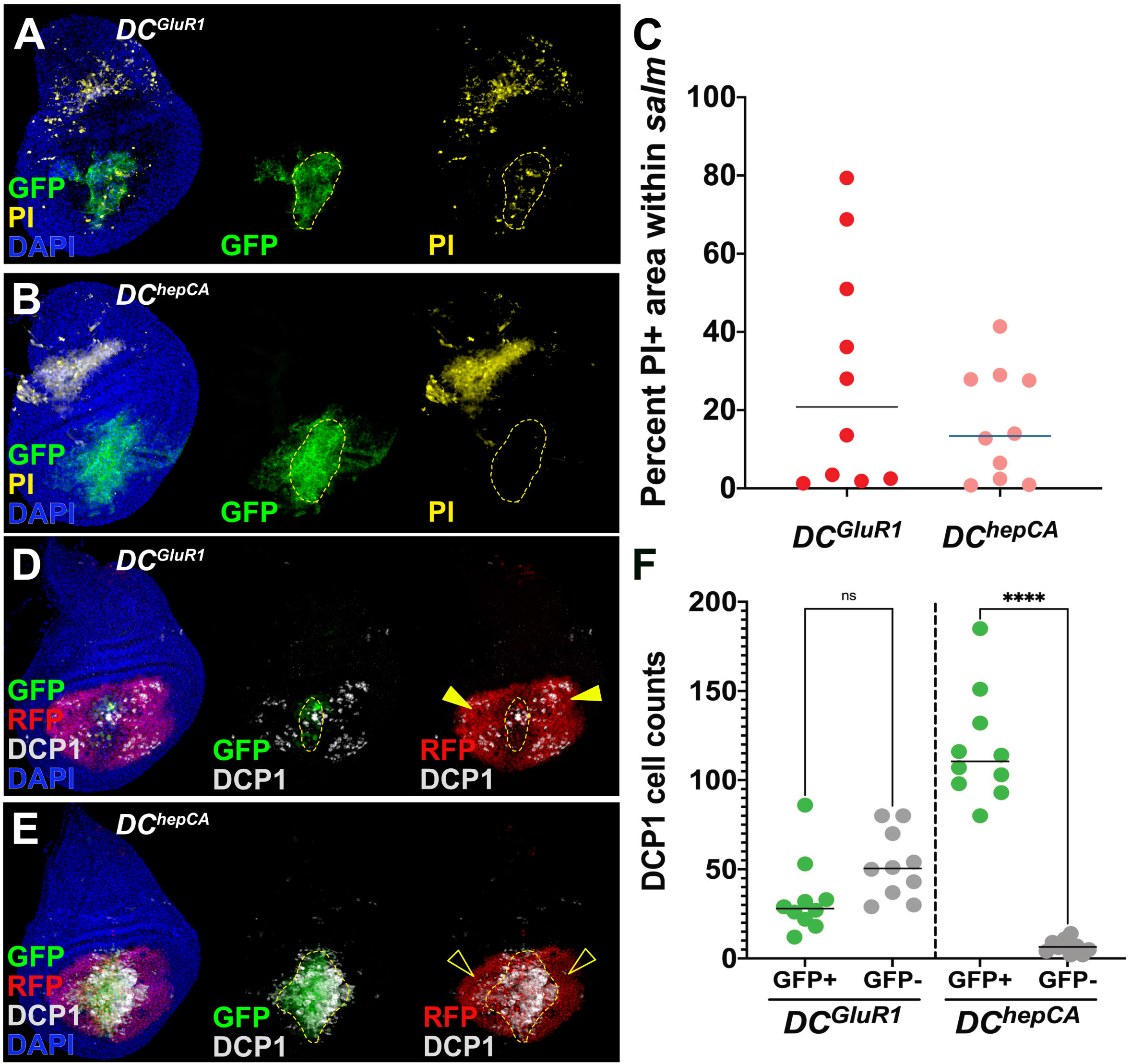
Quantification of cell death in GluRI ablated discs. (**A-B**) Wing discs bearing a *lexAop-GFP* transgene and ablated with *DC^GluR1^* (**A**) and *DC^hepCA^* (**B**) and stained for PI (yellow). DAPI, blue. PI staining in the notum is due to physical manipulation of the wing discs and acts as an internal control. (**C**) Quantification of the PI signal. The *lexAop-GFP* was used to outline the ablated tissue for quantification, as seen in (A) and (B). n = 10 discs for each treatment. (**D-E**) Wing imaginal discs bearing a *DR^WNT^-GFP ; UAS-RFP* transgene, ablated with *DC^GluR1^* (**D**) and *DC^hepCA^* (**E**), and stained for DCP1, gray. DAPI, blue. Solid arrowheads in (D) highlight GFP negative, DCP1 positive cells. Open arrowheads in (E) indicate a lack of GFP negative, DCP1 positive cells in *DC^hepCA^* ablated discs. (**F**). Quantification of DCP1 cells at the wound edge (GFP+) vs. the lateral wing pouch (GFP-). ns: not significant. P**** < 0.0001. Data analyzed with a Kruskal-Wallis test and a Tukey’s multiple comparisons test. n = 10 discs for each treatment.

**Figure S3. Related to Figure 3.**
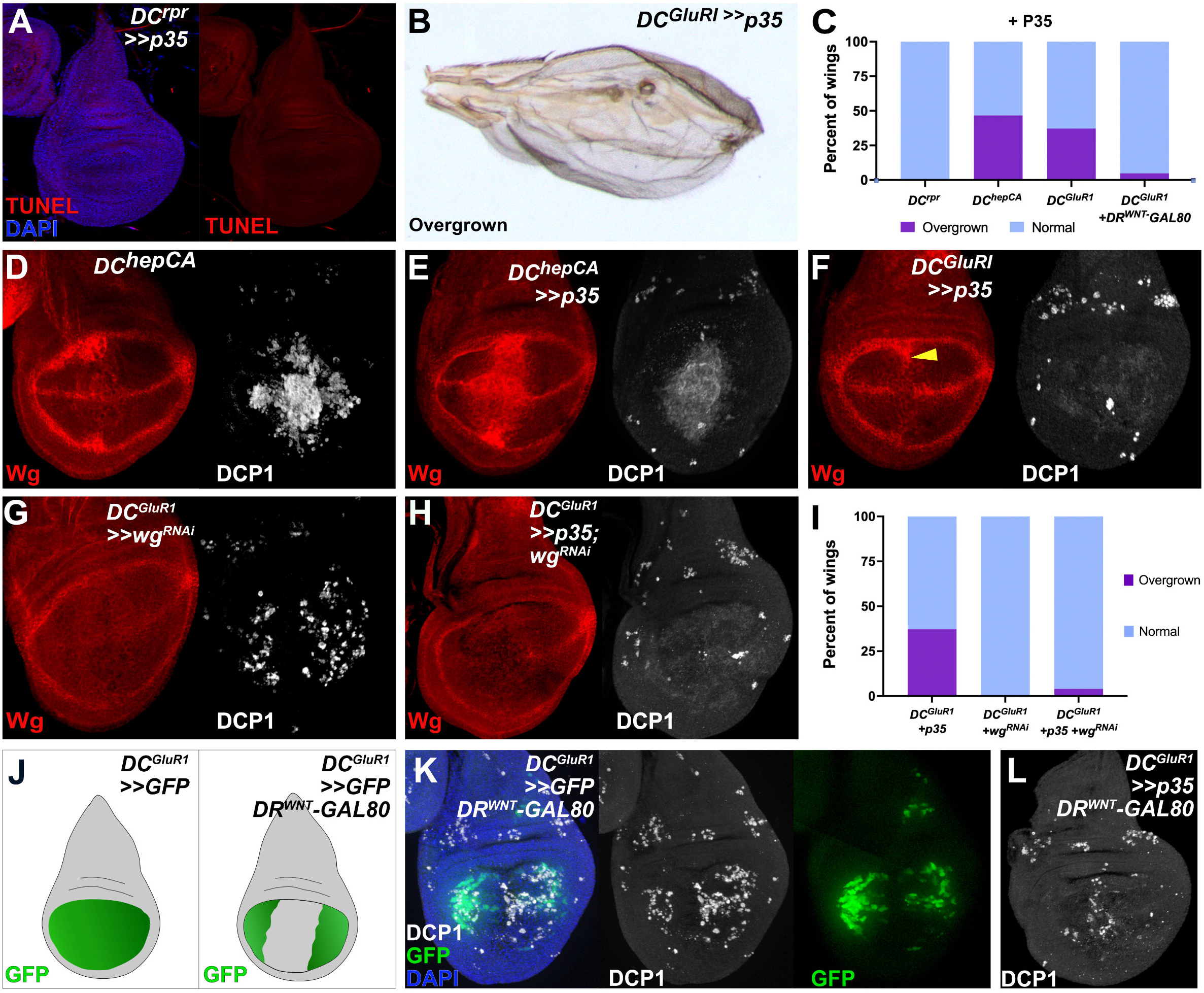
Blocking apoptosis with P35 leads to wing overgrowth phenotypes. (**A**) A wing disc ablated with *DC^rpr^>>p35* and labeled with TUNEL (red). DAPI, blue. The lack of signal demonstrates the ability of P35 to inhibit cell death. (**B**) Adult wings showing the overgrown phenotype accompanied by blistering and ectopic vein tissue observed following *DC^GluR1^>>p35* ablation. (**C**) Quantification of overgrown wing phenotype frequency following ablation with *DC^rpr^>>p35* (n = 89), *DC^hepCA^>>p35* (n = 30)*, DC^GluR1^>>p35* (n = 43), and *DC^GluR1^ DR^Wnt^-GAL80 >>p35* (104). No overgrown wings are observed in *DC^rpr^>>p35*. (**D-F**) Wing imaginal discs stained for Wg (red) and DCP1 (gray) following ablation with *DC^hepCA^* (**D**), *DC^hepCA^>>p35* (**E**) or *DC^GluR1^>>p35* (**F**). Ectopic Wg overlaps with the diffuse DCP1 signal in *DC^hepCA^>>p35* discs, while in *DC^GluR1^>>p35* discs ectopic Wg is only observed at the wound edge (arrowhead) and not in the diffuse DCP1 signal in the rest of the pouch. (**G-H**) Wing imaginal discs ablated with *DC^GluR1^>>wg^RNAi^* (**G**) *DC^GluR1^>>p35; wg^RNAi^* (**H**) and stained for Wg (red) and DCP1 (gray). (**I**) Quantification of overgrown wing phenotype frequency following ablation with *DC^GluR1^* (n = 43)*, DC^GluR1^>>^wgRNAi^* (n = 62), and *DC^GluR1^>>p35; wg^RNAi^* (n = 46). (**J**) Schematic of wing discs bearing *DC^GluR1^>>GFP* with and without *DR^WNT^-GAL80*, demonstrating the ability of *DR^WNT^-GAL80* to limit *UAS-GFP* expression to the lateral pouch. (**K**) A wing disc ablated by *DC^GluR1^>>GFP, DR^Wnt^-GAL80* and stained for DCP1 (gray). DAPI, blue. The GFP signal overlaps with the NiA cells but not the wound edge apoptosis. (**L**) A wing disc ablated by *DC^GluR1^>>p35, DR^Wnt^-GAL80* and stained for DCP1 (gray). DAPI, blue. The wound edge DCP1 is still strong and punctate while the NiA cells are no longer visible.

**Figure S4. Related to Figure 5.**
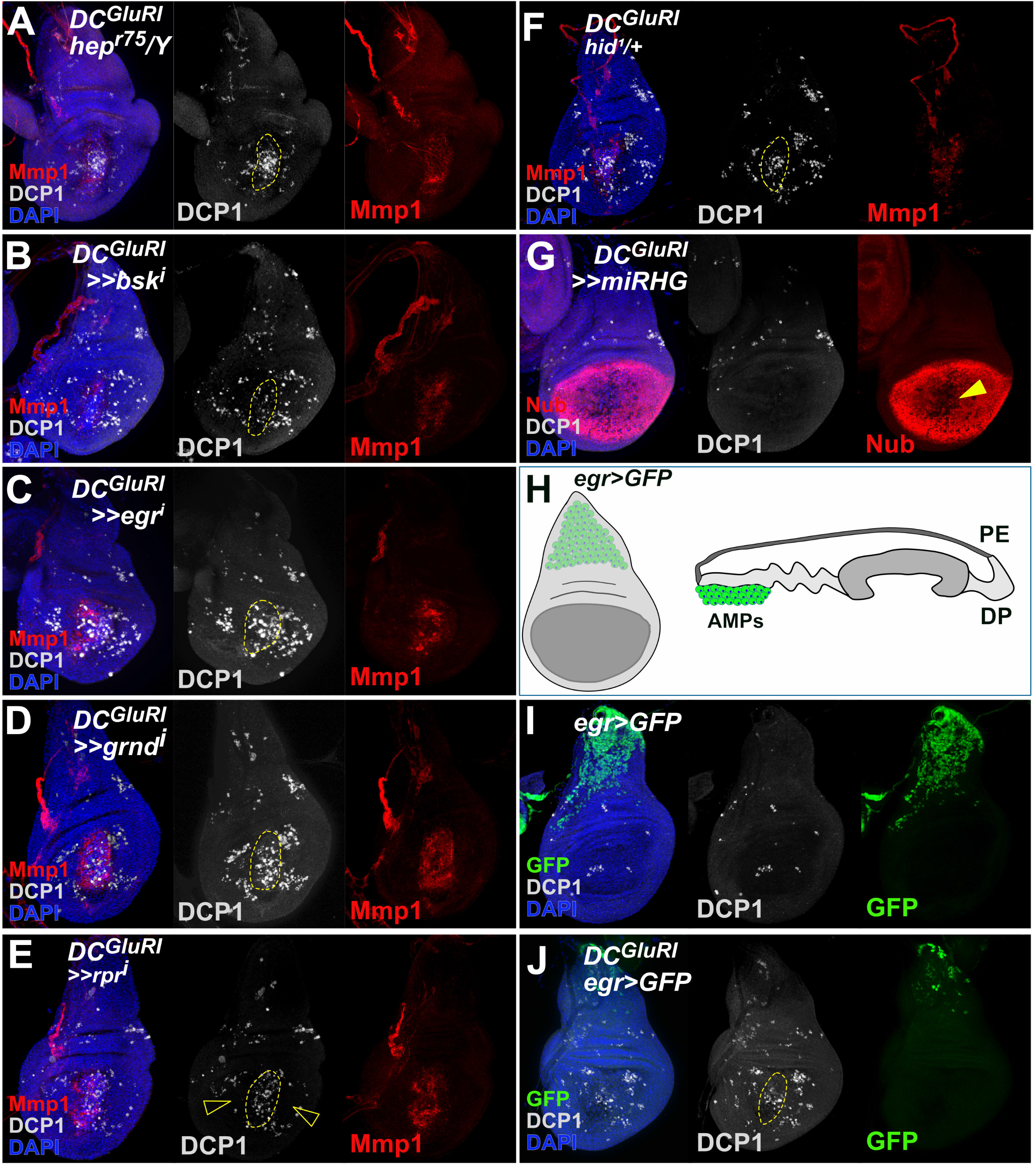
Manipulation of JNK pathway elements in *DC^GluRI^* ablated discs. (**A**) A wing disc ablated with *DC^GluR1^* in a *hep^r75^* hemizygous background and stained for DCP1 (gray) and Mmp1 (red). DAPI, blue. Wound edge apoptosis and NiA cells are still observed in this background, although are overall reduced. As in other panels, the dashed line outlines the ablated *salm* domain, indicated by the change in DAPI appearance. (**B**) A wing disc ablated with *DC^GluR1^>>bsk^RNAi^* and stained for DCP1 (gray) and Mmp1 (red), DAPI, blue. Wound edge apoptotic cells are reduced following *bsk* knockdown, while NiA cells remain unaffected. (**C**) A wing disc ablated with *DC^GluR1^>>egr^RNAi^* stained for DCP1 (gray) and Mmp1 (red). DAPI, blue. Both wound edge apoptotic cells and NiA cells remain unaffected by *egr* knockdown. (**D**) A wing disc ablated with *DC^GluR1^>>grnd^RNAi^* and stained for DCP1 (gray) and Mmp1 (red). DAPI, blue. As in (**C**), both wound edge apoptotic cells and NiA cells remain unaffected following *grnd* knockdown. (**E**) A wing disc ablated with *DC^GluR1^>>rpr^RNAi^* and stained for DCP1 (gray) and Mmp1 (red). DAPI, blue. Knockdown of *rpr* leads to a mild reduction of both wound edge apoptotic cells and NiA cells (open arrowheads). (**F**) A wing disc ablated with *DC^GluR1^* in a *hid^1^* heterozygous background, and stained for DCP1 (gray) and Mmp1 (red), DAPI, blue. Both wound edge apoptotic cells and NiA cells are unaffected in this background. (**G**) A wing imaginal disc ablated with *DC^GluR1^>>miRHG* and stained for DCP1 (gray) and Nub (red). DAPI, blue. Tissue loss indicated by loss of Nub (arrowhead) is still observed following ablation with *DC^GluR1^* despite inhibition of apoptosis by knockdown of *rpr, hid,* and *grim*. (**H**) Schematic of the *egr-Gal4 ; UAS-GFP* reporter showing activity in the adult muscle precursor cells (AMPs) just below the notum. (**I-J**) Wing imaginal discs bearing an *egr* reporter (*egr-GAL4; UAS-GFP*, green) in an undamaged heat shocked disc (**I**) and following ablation with a version of *DC^GluR1^* that lacks the *DVE>>GAL4* (**J**). DAPI, blue. Reporter activity is not observed in the pouch with or without ablation

**Figure S5. Related to Figure 6.**
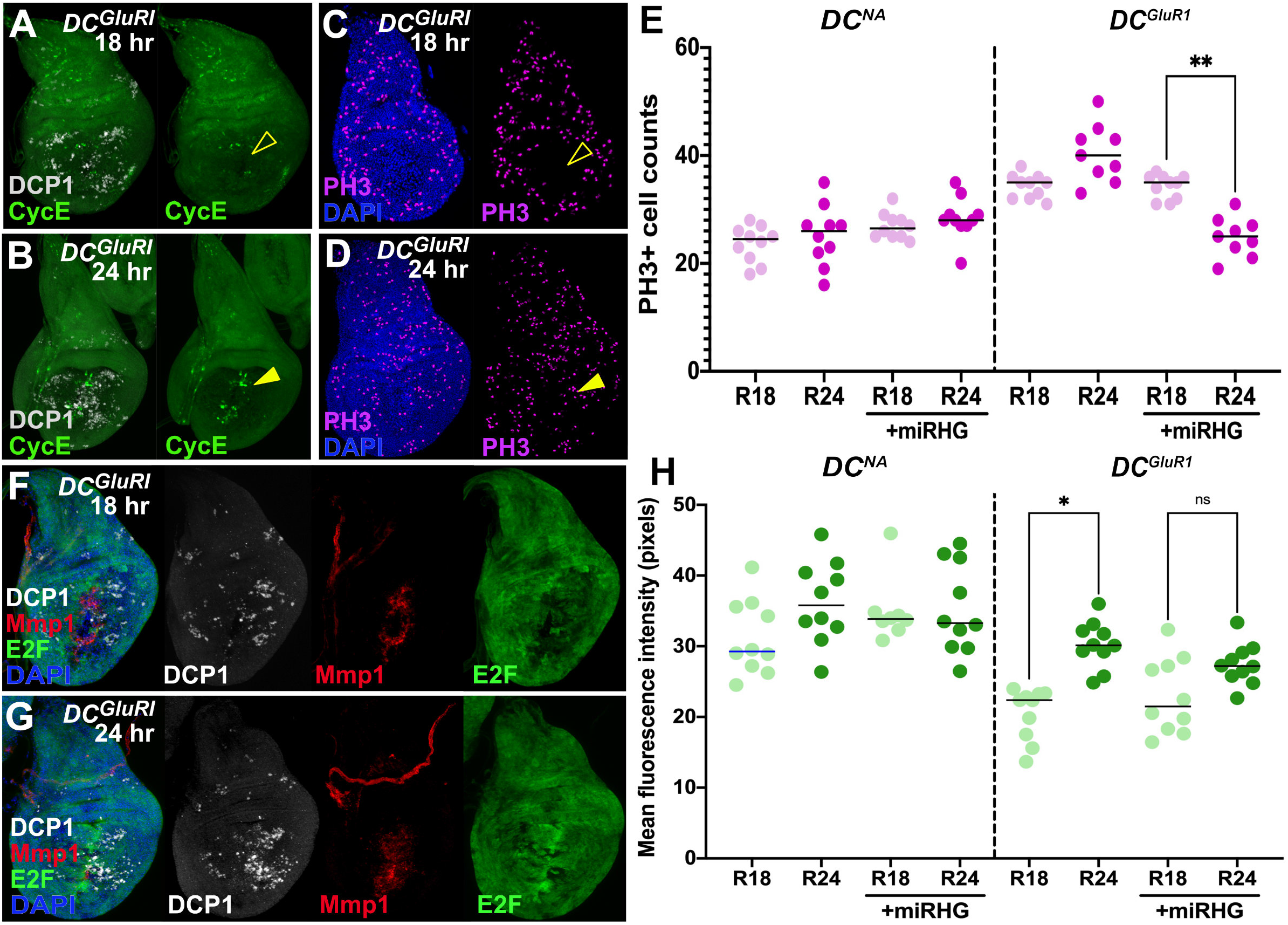
Quantification of proliferation with NiA manipulation. (**A-B)** Wing imaginal discs bearing the *CycE-GFP* reporter (green) at 18 hr (**A**) and 24 hr (**B**) of recovery following ablation by *DC^GluR1^* and stained for DCP1 (gray). DAPI, blue. The open arrowhead in (A) indicates a lack of reporter activity in the *salm* domain, while the arrowhead in (B) highlights stronger reporter activity at 24 hr. (**C-D**) Wing imaginal discs after 18 hr (**C**) and 24 hr (**D**) of recovery following *DC^GluR1^* ablation, and stained for DCP1 (gray), PH3 (magenta), and DAPI. The open arrowhead in (C) indicates a lack of PH3 in the *salm* domain, while the arrowhead in (D) highlights increased PH3 at 24 hr. (**E**) Quantification of PH3+ cells in *DC^NA^* disc and in *DC^GluR1^* ablated discs, both in the presence and absence of *miRHG* and after 18 hr and 24 hr after the heat shock/recovery. P** < 0.005. Data was analyzed via a Kruskal-Wallis test and a Tukey’s multiple comparisons test. n = 10 discs for each treatment. (**F-G**) Wing discs bearing the *PCNA-GFP* reporter at 18 hr (**F**) and 24 hr (**G**) of recovery following *DC^GluR1^* ablation, and stained for DCP1 (gray) and Mmp1 (red). DAPI, blue. (**H**) Quantification of the mean fluorescence intensity of the *PCNA-GFP* signal at 18 hr and 24 hr of recovery in the presence and absence of *miRHG*, following *DC^GluR1^* ablation. ns. P* < 0.0445. Data was analyzed via a Kruskal-Wallis test and a Tukey’s multiple comparisons test. n = 10 discs for each treatment.

## Notes

### Competing Interest Statement

The authors have declared no competing interest.

### Summary of Updates

Additional experiments, quantification and analyses performed.

