## Supplemental Methods for "Necrosis-induced apoptosis promotes regeneration in *Drosophila* wing imaginal discs"

### Figure Genotypes

#### DUAL Control genotypes

$DC^{NA}$
*hs-flp / + ; hs-p65 / + ; salm-LexADBD , DVE>>GAL4 / +*
$DC^{GluR1}$
*hs-flp / + ; lexAop-GluR1*$^{LC}$ *, hs-p65 / + ; salm-LexADBD , DVE>>GAL4 / +*
$DC^{GluR1;NoGAL4}$
*hs-flp / + ; lexAop-GluR1*$^{LC}$ *, hs-p65 / + ; salm-LexADBD / +*
$DC^{rpr}$
*hs-flp / + ; lexAop-rpr , hs-p65 / + ; salm-LexADBD , DVE>>GAL4 / +*
$DC^{hepCA}$
*hs-flp / + ; lexAop-hepCA , hs-p65 / + ; salm-LexADBD , DVE>>GAL4 / +*

#### Figure 1

(**C**, **H**, **I**) *hs-flp / + ; hs-p65 / + ; salm-LexADBD , DVE>>GAL4 / UAS-y*$^{RNAi}$
(**D**, **H**, **I**) *hs-flp / + ; lexAop-GluR1*$^{LC}$ *, hs-p65 / + ; salm-LexADBD , DVE>>GAL4 / UAS-y*$^{RNAi}$
(**E**, **H**, **I**) *hs-flp / + ; lexAop-rpr , hs-p65 / + ; salm-LexADBD , DVE>>GAL4 / UAS-y*$^{RNAi}$
(**F**, **H**, **I**) *hs-flp / + ; lexAop-hepCA , hs-p65 / + ; salm-LexADBD , DVE>>GAL4 / UAS-y*$^{RNAi}$

#### Figure 2

(**A**, **D**, **K**) *hs-flp / + ; lexAop-GluR1*$^{LC}$ *, hs-p65 / + ; salm-LexADBD , DVE>>GAL4 / UAS-y*$^{RNAi}$
(**B**, **F**) *hs-flp / + ; lexAop-hepCA , hs-p65 / + ; salm-LexADBD , DVE>>GAL4 / UAS-y*$^{RNAi}$
(**C**) *hs-flp / + ; hs-p65 / + ; salm-LexADBD , DVE>>GAL4 / UAS-y*$^{RNAi}$
(**E**, **L**) *hs-flp / + ; lexAop-rpr , hs-p65 / + ; salm-LexADBD , DVE>>GAL4 / UAS-y*$^{RNAi}$
(**G**-**H**) *hs-flp / + ; lexAop-GluR1*$^{LC}$ *, hs-p65 / lexAop-GFP ; salm-LexADBD , DVE>>GAL4 / UAS-RFP*
(**I**-**J**) *hs-flp / + ; lexAop-hepCA , hs-p65 / lexAop-GFP ; salm-LexADBD , DVE>>GAL4 / UAS-RFP*

#### Figure 3

(**A**, **D**, **I**, **K’**, **M**) *hs-flp / + ; lexAop-rpr , hs-p65 / + ; salm-LexADBD , DVE>>GAL4 / UAS-p35*
(**B**, **F**, **G**, **H**, **J**, **L’**, **M**) *hs-flp / + ; lexAop-GluR1*$^{LC}$ *, hs-p65 / + ; salm-LexADBD , DVE>>GAL4 / UAS-p35*
(**C**, **I**, **K**, **M**) *hs-flp / + ; lexAop-rpr, hs-p65 / + ; salm-LexADBD , DVE>>GAL4 / UAS-y*$^{RNAi}$
(**E**, **J**, **L**, **M**) *hs-flp / + ; lexAop-GluR1*$^{LC}$ *, hs-p65 / + ; salm-LexADBD , DVE>>GAL4 / UAS-y*$^{RNAi}$

#### Figure 4

(**A**-**F**) *hs-flp / + ; lexAop-GluR1*$^{LC}$ *, hs-p65 / DR*$^{WNT}$*-GFP ; salm-LexADBD , DVE>>GAL4 / +*
(**G**) *hs-flp / + ; lexAop-GluR1*$^{LC}$ *, hs-p65 / + ; salm-LexADBD , DVE>>GAL4 / UAS-y*$^{RNAi}$

**Figure 5**

(**A**) hs-flp / + ; lexAop-GluR1$^{LC}$ , hs-p65 / AP-1-GFP ; salm-LexADBD , DVE>>GAL4 / +
(**B**) hs-flp / + ; lexAop-GluR1$^{LC}$ , hs-p65 / + ; salm-LexADBD , DVE>>GAL4 / puc$^{A251}$
(**C**-**D**) hs-flp / + ; lexAop-GluR1$^{LC}$ , hs-p65 / + ; salm-LexADBD , DVE>>GAL4 / UAS-y$^{RNAi}$
(**E**) hs-flp / + ; lexAop-GluR1$^{LC}$ , hs-p65 / + ; salm-LexADBD , DVE>>GAL4 / UAS-hep$^{RNAi}$
(**F**) hs-flp / + ; lexAop-GluR1$^{LC}$ , hs-p65 / + ; salm-LexADBD , DVE>>GAL4 / UAS-bsk$^{DN}$
(**G**) hs-flp / + ; lexAop-GluR1$^{LC}$ , hs-p65 / + ; salm-LexADBD , DVE>>GAL4 / UAS-mkk4$^{RNAi}$
(**H**) hs-flp / + ; lexAop-GluR1$^{LC}$ , hs-p65 / UAS-miRHG ; salm-LexADBD , DVE>>GAL4 / +

**Figure 6**

(**A**, **B**) hs-flp / + ; lexAop-GluR1$^{LC}$ , hs-p65 / + ; salm-LexADBD , DVE>>GAL4 / PCNA-GFP
(**C**) hs-flp / + ; lexAop-GluR1$^{LC}$ , hs-p65 / UAS-miRHG ; salm-LexADBD , DVE>>GAL4 / PCNA-GFP
(**D**, **F**, **G**) hs-flp / + ; lexAop-GluR1$^{LC}$ / UAS-miRHG ; salm-LexADBD , DVE>>GAL4 / DR$^{WNT}$-GAL80
(**E**, **G**) hs-flp / + ; lexAop-rpr , hs-p65 / + ; salm-LexADBD , DVE>>GAL4 / UAS-y$^{RNAi}$
hs-flp / + ; lexAop-rpr , hs-p65 / UAS-miRHG ; salm-LexADBD , DVE>>GAL4 / +
(**F**, **G**) hs-flp / + ; lexAop-GluR1$^{LC}$ , hs-p65 / + ; salm-LexADBD , DVE>>GAL4 / UAS-y$^{RNAi}$
hs-flp / + ; lexAop-GluR1$^{LC}$ , hs-p65 / UAS-miRHG ; salm-LexADBD , DVE>>GAL4 / +

**Supplemental Figures**

**Figure S1**

(**B**) hs-flp / + ; hs-p65 / lexAop-GFP ; salm-LexADBD , DVE>>GAL4 / UAS-RFP
(**C**) hs-flp / + ; lexAop-GluR1$^{LC}$ , hs-p65 / + ; salm-LexADBD , DVE>>GAL4 / +
(**D**) hs-flp / + ; hs-p65 / + ; salm-LexADBD , DVE>>GAL4 / +
(**E**) hs-flp / + ; lexAop-GluR1$^{LC}$ / + ; salm-LexADBD , DVE>>GAL4 / +
(**F**) hs-flp / + ; lexAop-GluR1$^{LC}$ , hs-p65 / + ; DVE>>GAL4 / +

**Figure S2**

(**A**, **C**) hs-flp / + ; lexAop-GluR1$^{LC}$ , hs-p65 / lexAop-GFP ; salm-LexADBD , DVE>>GAL4 / +
(**B**, **C**) hs-flp / + ; lexAop-hepCA , hs-p65 / lexAop-GFP ; salm-LexADBD , DVE>>GAL4 / +
(**D**, **F**) hs-flp / + ; lexAop-GluR1$^{LC}$ , hs-p65 / DR$^{WNT}$-GFP ; salm-LexADBD , DVE>>GAL4 / UAS-mCherry
(**E**, **F**) hs-flp / + ; lexAop-hepCA , hs-p65 / DR$^{WNT}$-GFP ; salm-LexADBD , DVE>>GAL4 / UAS-mCherry

**Figure S3**

(**A**, **C**) *hs-flp / + ; lexAop-rpr , hs-p65 / + ; R85E08-DBD , DVE>>GAL4 / UAS-p35*
(**B**, **C**, **F**, **I**) *hs-flp / + ; lexAop-GluR1*$^{LC}$ *, hs-p65 / + ; R85E08-DBD , DVE>>GAL4 / UAS-p35*
(**C**, **E**) *hs-flp / + ; lexAop-hepCA , hs-p65 / + ; salm-LexADBD , DVE>>GAL4 / UAS-p35*
(**C**, **L**) *hs-flp / + ; lexAop-GluR1*$^{LC}$ *, hs-p65 / UAS-p35 ; salm-LexADBD , DVE>>GAL4 / DR*$^{WNT}$*-GAL80*
(**D**) *hs-flp / + ; lexAop-hepCA , hs-p65 / + ; salm-LexADBD , DVE>>GAL4 / UAS-y*$^{RNAi}$
(**G**, **I**) *hs-flp / + ; lexAop-GluR1*$^{LC}$ */ + ; salm-LexADBD , DVE>>GAL4 / UAS-wg*$^{RNAi}$
(**H**, **I**) *hs-flp / + ; lexAop-GluR1*$^{LC}$ */ UAS-p35 ; salm-LexADBD , DVE>>GAL4 / UAS-wg*$^{RNAi}$
(**K**) *hs-flp / + ; lexAop-GluR1*$^{LC}$ *, hs-p65 / UAS-GFP ; salm-LexADBD , DVE>>GAL4 / DR*$^{WNT}$*-GAL80*

**Figure S4**
(**A**) *hep*$^{r75}$ */ Y ; lexAop-GluR1*$^{LC}$ *, hs-p65 / + ; salm-LexADBD , DVE>>GAL4 / +*
(**B**) *hs-flp / + ; lexAop-GluR1*$^{LC}$ *, hs-p65 / + ; salm-LexADBD , DVE>>GAL4 / UAS-bsk*$^{RNAi}$
(**C**) *hs-flp / + ; lexAop-GluR1*$^{LC}$ *, hs-p65 / + ; salm-LexADBD , DVE>>GAL4 / UAS-egr*$^{RNAi}$
(**D**) *hs-flp / + ; lexAop-GluR1*$^{LC}$ *, hs-p65 / + ; salm-LexADBD , DVE>>GAL4 / UAS-grnd*$^{RNAi}$
(**E**) *hs-flp / + ; lexAop-GluR1*$^{LC}$ *, hs-p65 / + ; salm-LexADBD , DVE>>GAL4 / UAS-rpr*$^{RNAi}$
(**F**) *hs-flp / + ; lexAop-GluR1*$^{LC}$ *, hs-p65 / + ; salm-LexADBD , DVE>>GAL4 / hid*$^{1}$
(**G**) *hs-flp / + ; lexAop-GluR1*$^{LC}$ *, hs-p65 / UAS-miRHG ; salm-LexADBD , DVE>>GAL4 / +*
(**I**, **J**) *hs-flp / + ; lexAop-GluR1*$^{LC}$ *, hs-p65 / UAS-GFP ; salm-LexADBD / egr-GAL4*

**Figure S5**
(**A**, **B**) *hs-flp / + ; lexAop-GluR1*$^{LC}$ *, hs-p65 / + ; salm-LexADBD , DVE>>GAL4 / CycE-disc-GFP*
(**C**, **D, E**) *hs-flp / + ; lexAop-GluR1*$^{LC}$ *, hs-p65 / + ; salm-LexADBD , DVE>>GAL4 / UAS-y*$^{RNAi}$
(**E**) *hs-flp / + ; hs-p65 / + ; salm-LexADBD , DVE>>GAL4 / UAS-yRNAi*
*hs-flp / + ; hs-p65 / UAS-miRHG ; salm-LexADBD , DVE>>Gal4 / +*
*hs-flp / + ; lexAop-GluR1*$^{LC}$ *, hs-p65 / UAS-miRHG ; salm-LexADBD , DVE>>GAL4 / +*
(**F**, **G**, **H**) *hs-flp / + ; lexAop-GluR1*$^{LC}$ *, hs-p65 / + ; salm-LexADBD , DVE>>GAL4 / PCNA-GFP*
(**H**) *hs-flp / + ; lexAop-GluR1*$^{LC}$ *, hs-p65 / UAS-miRHG ; salm-LexADBD , DVE>>GAL4 / PCNA-GFP*
